# CYClones: A highly powered, fully genotyped, 8-parent yeast mapping population

**DOI:** 10.1101/2025.10.15.682626

**Authors:** Gareth A. Cromie, Russell S. Lo, Trey S. Morgan, Anne E. Clark, Julee Ashmead, Martin S. Timour, Amy Sirr, Joshua M. Akey, Aimée M. Dudley

## Abstract

The budding yeast *Saccharomyces cerevisiae* is a remarkably adaptable organism that thrives in diverse environments. Global sequencing of natural isolates has revealed extensive genetic diversity within the species. Here, we describe the construction and characterization of CYClones (Collaborative Yeast Cross clones), a library of 11,392 segregants generated from a multiparent funnel cross of eight genetically diverse parental strains. To enable the genetic dissection of complex traits, we imputed whole-genome sequences for all segregants and show that CYClones captures a substantial fraction of the global genetic diversity of *S. cerevisiae*. Haplotype representation is well maintained, with each parental haplotype present at >5% frequency across >95% of the genome. Simulations demonstrate that CYClones has ≥95% power to detect variants with heritability as low as 0.36%, with mapping resolution often finer than the length of a single gene. In summary, CYClones is a powerful community resource for dissecting the genetic architecture of complex and quantitative traits, uncovering context-dependent mutational effects, and identifying causal variants underlying phenotypic diversity.

## Introduction

With its compact genome, deeply rooted connections to world culture, and prominence as a powerful model organism, the budding yeast *Saccharomyces cerevisiae* served as an ideal candidate for early whole genome sequencing (WGS) efforts. A seven-year collaboration among over 100 laboratories sequenced the 12 million base pairs of the widely used laboratory strain S288c, making it the first eukaryote to have its genome fully sequenced [1, 2]. Over the last several decades, sequencing costs have plummeted, and a concerted effort within the yeast community has resulted in the DNA sequence characterization of large collections of natural variant strains that have been isolated from both human-associated activities and diverse wild ecological niches.

Evidence of the domestication of yeast currently dates to China over 7,000 years ago [3, 4]. The use of *S. cerevisiae* as a leavening agent, or fermenter, has since been adopted by communities across the globe, creating foods and beverages as diverse as the heritages which produced them. *S. cerevisiae* has, therefore, been exposed and adapted to many different conditions and environments. In order to capture the diversity imposed by these selective pressures, budding yeast strains have been isolated and sequenced from human-associated products such as wine, beer, sake, mead, olives, raw coffee beans and cacao, from natural sources including tree bark, soil, insects, and from both immunosuppressed and healthy human subjects [5–16]. In addition to expanding the known genetic and phenotypic diversity beyond the well-characterized lab strains, these collections have greatly expanded our understanding of the evolutionary history, origins, and population structure of *S. cerevisiae* [17–22].

The genetic variation present in natural populations represents a powerful tool for interrogating genotype-phenotype relationships. Among approaches to this question, genome-wide association studies (GWAS) capture the highest proportion of natural genetic variation through the use of large samples drawn directly from the full population [23]. However, population structure can lead to spurious associations in GWAS studies and, until recently, the complex and heterogenous population structures inherent in the limited number of strains sequenced have made GWAS very challenging in yeast [24]. WGS of 1,011 natural variant yeast strains has helped ameliorate this problem, and GWAS has now been carried out in yeast, detecting the effects of many genetic variants across numerous conditions [9]. However, as in other organisms, despite capturing a large portion of the known genetic diversity within the species, this set of strains remains underpowered to detect the impact of rare variants with a frequency lower than 5% within the population. In addition, population structure constrains allele assortment, so that certain combinations of alleles are absent or severely under-represented among natural strains.

Mapping populations generated through crossing, and analyzed by linkage analysis, represent an alternative approach to interrogating genotype-phenotype relationships. These populations can be generated from two (pairwise) or more (multiparent) founders. In addition to GWAS studies, natural variant yeast collections have been leveraged as a resource for multiparent quantitative trait locus (QTL) mapping experiments [22, 25–28]. Relative to pairwise approaches, multiparent study designs enable a larger and more diverse set of alleles to be analyzed in a single experiment. In addition, any rare (in the natural population) alleles captured in a multiparent cross will be present in the mapping population at frequencies similar to common alleles captured, resulting in approximately equal power to detect the phenotypic effects of rare and common variants. Multiparent crosses also eliminate the confounding effects of population structure that complicates GWAS. However, many designs impose other forms of population structure that still limit the mixing of rare alleles from different genetic backgrounds.

We set out to develop a library of yeast strains that captures a substantial fraction of genomic variation in yeast and that avoids the confounding effects of population structure. Multiparent “funnel cross” breeding schemes generate populations that harbor all combinations of parental alleles and have been proposed as an efficient means of accomplishing these goals [29, 30]. Therefore, we constructed CYClones (<u>C</u>ollaborative <u>Y</u>east <u>C</u>ross clones), a set of 11,392 fully genotyped recombinant strains generated from eight parental yeast strains using a funnel cross design. CYClones complements complex crosses in other model organisms (generally referred to as <u>M</u>ulti-parent <u>A</u>dvanced <u>G</u>eneration Inter-<u>C</u>ross lines), such as MAGIC [31, 32] in *Arabidopsis*, Flyland [33] in *Drosophila*, and the mouse collaborative cross [34]. Indeed, our eight-parent funnel cross in the budding yeast, *Saccharomyces cerevisiae* (Fig 1) was inspired by the Mouse Collaborative Cross [34], which aimed to generate 1000 recombinant inbred lines derived from eight parental mouse strains. However, since that project was hampered by exceptionally high (95%) extinction of the recombinant inbred lines during the final inbreeding process [34], a large, highly powered multiparent mapping population generated via such a design has yet to be examined.

**Fig 1.**
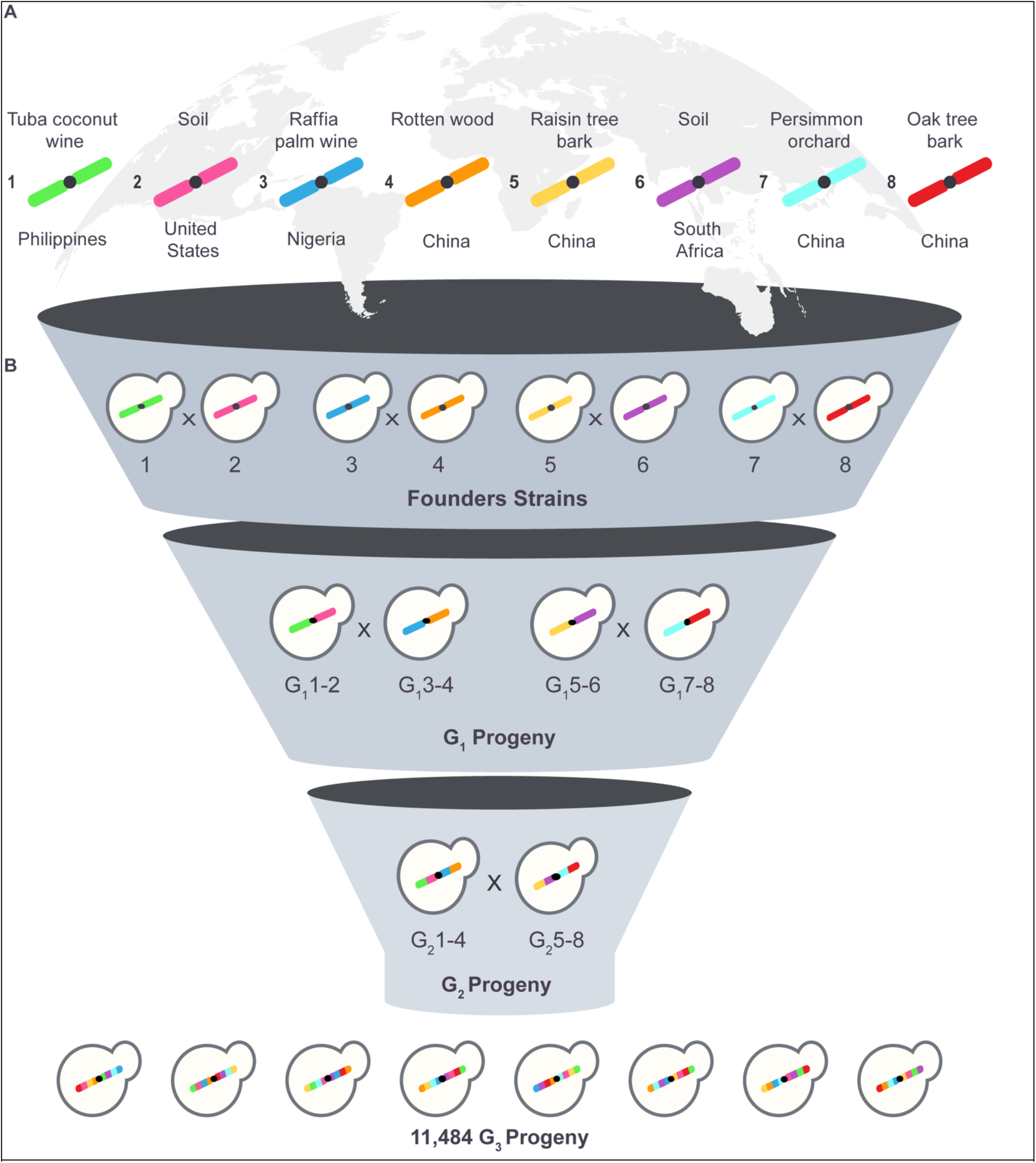
Generation of the mapping population. (A) Eight haploid strains from diverse geographical and environmental sources, harboring a substantial proportion of the rare and common alleles present in the global *S. cerevisiae* population, were chosen to generate a multi-parent mapping population. (B) The mapping population was constructed using a funnel cross design. Each founder strain was mated pairwise to produce pools of haploid progeny. Two rounds of reciprocal crossing using pooled strains were then carried out to produce the final mapping population of ∼11,400 haploid strains, each a recombinant mosaic of the eight founder haplotypes.

## Materials and Methods

### Media and genetic manipulation of yeast

Unless noted, standard media and methods were used for the growth and manipulation of yeast [35]. The *S. cerevisiae* strains used in this study are listed in Table 1. Unless specified, yeast strains were grown in YPD medium (1% yeast extract, 2% bacto peptone, 2% glucose) with 2% agar for solid plates. Nourseothricin (NAT) was used at a final concentration of 100 µg/ml. Hygromycin (HYG) was used at a final concentration of 500 µg/ml.

**Table 1.** *S. cerevisiae* strains used in this study.

| Strain Name | Mating Type | Derived From | Geographic Origin | Reference | Genotype |
| --- | --- | --- | --- | --- | --- |
| YO1048 (NRRL Y-6297) | MATa/α | N. A | Philippines tuba (coconut sap wine) | Ludlow et al. [6] |  |
| YO302 (IL-01) | MATa/α | N. A | Cahokia, Illinois, USA soil sample at base of oak tree | Ludlow et al. [6] |  |
| YO303 (PW5) | MATa/α | N. A | Aba, Abia State, Nigeria, Africa Raphia palm wine | Ludlow et al. [6] |  |
| YO1560 (HN15) | MATa/α | N. A | Wuzhi Mountain, Hainan Island, China Rotten wood in primeval, tropical forest | Wang et al. [10] |  |
| YO1568 (FJ7) | MATa/α | N. A | Wuyi Mountain Fujian, China Bark of Hovenia acerba (Chinese raisin tree), subtropical primeval forest | Wang et al. [10] |  |
| YO1047 (Y-12638, CBS 2888) | MATa/α | N. A | South Africa, Africa Soil sample | Ludlow et al. [6] |  |
| YO1562 (BJ6) | MATa/α | N. A | Changping, Beijing, China Persimmon orchard | Wang et al. [10] |  |
| YO1567 (SX6) | MATa/α | N. A | Qinling Monutain, Shaanxi, China Bark of Quercus fabri (oak) tree, temperate primeval forest | Wang et al. [10] |  |
| YO2644 | MATa/α | YO1048 | Philippines tuba (coconut sap wine) | This study | *HO / hoΔ1198::hisG:natMX6:yeGFP::hisG* |
| YO2379 | MATa/α | YO302 | Cahokia, Illinois, USA soil sample at base of oak tree | This study | *HO / hoΔ1198::hisG:natMX6:yeGFP::hisG* |
| YO2409 | MATa/α | YO303 | Aba, Abia State, Nigeria, Africa Raphia palm wine | This study | *HO / hoΔ1198::hisG:natMX6:yeGFP::hisG* |
| YO2391 | MATa/α | YO1560 | Wuzhi Mountain, Hainan Island, China Rotten wood in primeval, tropical forest | This study | *HO / hoΔ1198::hisG:natMX6:yeGFP::hisG* |
| YO2394 | MATa/α | YO1568 | Wuyi Mountain Fujian, China Bark of Hovenia acerba (Chinese raisin tree), subtropical primeval forest | This study | *HO / hoΔ1198::hisG:natMX6:yeGFP::hisG* |
| YO2410 | MATa/α | YO1047 | South Africa, Africa Soil sample | This study | *HO / hoΔ1198::hisG:natMX6:yeGFP::hisG* |
| YO2412 | MATa/α | YO1562 | Changping, Beijing, China Persimmon orchard | This study | *HO / hoΔ1198::hisG:natMX6:yeGFP::hisG* |
| YO2382 | MATa/α | YO1567 | Qinling Monutain, Shaanxi, China Bark of Quercus fabri (oak) tree, temperate primeval forest | This study | *HO / hoΔ1198::hisG:natMX6:yeGFP::hisG* |
| YO2645 | MATa | YO2644 | Philippines tuba (coconut sap wine) | This study | *hoΔ1198::hisG:natMX6:yeGFP::hisG* |
| YO2443 | MATα | YO2379 | Cahokia, Illinois, USA soil sample at base of oak tree | This study | *hoΔ1198::hisG:natMX6:yeGFP::hisG* |
| YO2451 | MATa | YO2409 | Aba, Abia State, Nigeria, Africa Raphia palm wine | This study | *hoΔ1198::hisG:natMX6:yeGFP::hisG* |
| YO2424 | MATα | YO2391 | Wuzhi Mountain, Hainan Island, China Rotten wood in primeval, tropical forest | This study | *hoΔ1198::hisG:natMX6:yeGFP::hisG* |
| YO2414 | MATa | YO2394 | Wuyi Mountain Fujian, China Bark of Hovenia acerba (Chinese raisin tree), subtropical primeval forest | This study | *hoΔ1198::hisG:natMX6:yeGFP::hisG* |
| YO2458 | MATα | YO2410 | South Africa, Africa Soil sample | This study | *hoΔ1198::hisG:natMX6:yeGFP::hisG* |
| YO2460 | MATa | YO2412 | Changping, Beijing, China Persimmon orchard | This study | *hoΔ1198::hisG:natMX6:yeGFP::hisG* |
| YO2428 | MATα | YO2382 | Qinling Monutain, Shaanxi, China Bark of Quercus fabri (oak) tree, temperate primeval forest | This study | *hoΔ1198::hisG:natMX6:yeGFP::hisG* |
| YO2681 | MATa | YO2645 | Philippines tuba (coconut sap wine) | This study | *hoΔ1198::hisG* |
| YO2478 | MATα | YO2443 | Cahokia, Illinois, USA soil sample at base of oak tree | This study | *hoΔ1198::hisG* |
| YO2506 | MATa | YO2451 | Aba, Abia State, Nigeria, Africa Raphia palm wine | This study | *hoΔ1198::hisG* |
| YO2485 | MATα | YO2424 | Wuzhi Mountain, Hainan Island, China Rotten wood in primeval, tropical forest | This study | *hoΔ1198::hisG* |
| YO2495 | MATa | YO2414 | Wuyi Mountain Fujian, China Bark of Hovenia acerba (Chinese raisin tree), subtropical primeval forest | This study | *hoΔ1198::hisG* |
| YO2480 | MATα | YO2458 | South Africa, Africa Soil sample | This study | *hoΔ1198::hisG* |
| YO2487 | MATa | YO2460 | Changping, Beijing, China Persimmon orchard | This study | *hoΔ1198::hisG* |
| YO2492 | MATα | YO2428 | Qinling Monutain, Shaanxi, China Bark of Quercus fabri (oak) tree, temperate primeval forest | This study | *hoΔ1198::hisG* |
| YO2685 (founder1) | MATa | YO2681 | Philippines Tuba (coconut sap wine) | This study | *hoΔ1198::hisG, PHO87::hphMX6::ARS314* |
| YO2545 (founder2) | MATα | YO2478 | Cahokia, Illinois, USA Soil near oak tree | This study | *hoΔ1198::hisG, PHO87::natMX6::ARS314* |
| YO2563 (founder3) | MATa | YO2506 | Aba, Abia State, Nigeria, Africa Raphia palm wine | This study | *hoΔ1198::hisG, PHO87::hphMX6::ARS314* |
| YO2552 (founder4) | MATα | YO2485 | Wuzhi Mountain, Hainan Island, China Rotten wood in primeval, tropical forest | This study | *hoΔ1198::hisG, PHO87::natMX6::ARS314* |
| YO2633 (founder5) | MATa | YO2495 | Wuyi Mountain Fujian, China Bark of Hovenia acerba (Chinese raisin tree), subtropical primeval forest | This study | *hoΔ1198::hisG, PHO87::hphMX6::ARS314* |
| YO2547 (founder6) | MATα | YO2480 | South Africa, Africa Soil sample | This study | *hoΔ1198::hisG, PHO87::natMX6::ARS314* |
| YO2553 (founder7) | MATa | YO2487 | Changping, Beijing, China Persimmon orchard | This study | *hoΔ1198::hisG, PHO87::hphMX6::ARS314* |
| YO2588 (founder8) | MATα | YO2492 | Qinling Monutain, Shaanxi, China Bark of Quercus fabri (oak) tree, temperate primeval forest | This study | *hoΔ1198::hisG, PHO87::natMX6::ARS314* |

### Founder whole genome sequencing, assessment of global yeast variation captured, and phylogenetic tree construction

The genome sequences of each of the eight haploid founder strains used to construct the mapping population were determined as follows. Genomic DNA was isolated using a YeaStar Genomic DNA Extraction Kit (Zymo Research). DNA sequencing libraries were prepared using the Paired-End Sequencing Kit (Illumina) following the manufacturer’s instructions. Sequencing reads were generated on an Illumina NextSeq 500, with 38 × 37 bp paired-end reads for all strains, except YO2685 for which 43 x 32 bp paired-end reads were used. Adaptor sequences were trimmed using Cutadapt (v1.7.1) [36]. Sequences were then aligned to the S288c reference genome (R64-1-1) using BWA (v7.9) [37], using the “aln” option with quality trimming (threshold of Phred = 20) and allowing a maximum of six mismatches. PCR duplicates were removed using the SAMtools (v0.1.19) [38] “rmdup” command. BCFtools (v1.10.2) [38] was then used to generate a variant call file (vcf) for each parental strain, using the “mpileup” command with the parameter “-d 10000” and the “call” command with the “-c” flag (consensus caller).

The VCF files were then assessed at all biallelic SNV (single nucleotide variant) sites with <5% missing data identified in a genomic analysis of the global yeast population [9]. The proportion of global sites for which we identified both alleles among our founder strains was then calculated for a range of minor allele frequency cutoffs in the global dataset. For phylogenetic tree construction, the set of sites was then further reduced to only those with quality scores >1000 in the published global gVCF file [9]. The haploid founder strains were added to this dataset of global yeast strains, encoded as homozygous reference, homozygous alternative, or missing data at each site. A neighbor-joining tree was then generated using the dissimilarity distance, as described in Peter et al. [9].

### Identification of SNVs among the founder strains

The sequencing reads for each founder strain were also used to generate a *de novo* genome assembly for that strain, and this assembly was compared to the reference genome to identify SNVs. First, reads were error-corrected with SGA (v0.9.9) [39] using the “preprocess” command with the parameters “-m 25 -p 1 -q 10 -- permute-ambiguous” and were then corrected using the “correct” command with the parameter “-k 25”. The corrected reads were assembled into contigs using IDBA (v1.0.12) [40] with the parameters --mink 19 --maxk 35 --step 2 --seed_kmer 23”. The final contigs were aligned to the reference genome using the Mummer (v 3.23) “nucmer” command with the parameters “-c 100 -l 10 -b 200” and the alignments were then filtered using the “delta-filter” command and the parameters “-r -q”. Finally, bases were called in the aligned regions using the “show-SNVs” command.

A set of 276,774 high-quality SNVs segregating in the mapping population was then identified by combining the VCF files from each founder strain with the basecalls from the *de novo* assembly pipeline (File S1: https://doi.org/10.5281/zenodo.17362654). Only biallelic SNVs with basecalls in all founder strains, consistent in both the VCF and *de novo* files for each strain, were accepted. In each VCF file, a basecall quality cutoff of greater than or equal to 50 was also applied and, for non-reference calls, the homozygous model for that basecall had to be more probable than the homozygous reference or heterozygous models.

### Constructing the mapping population

To prevent mating type switching, and thus maintaining stable haploid strains at each level of the cross, *hoΔ0* haploid founder strains were generated from the original diploid strains (Table 1) as follows. First, in each diploid strain, one copy of the gene encoding the HO endonuclease was deleted by integration of an HO-hisG-NatMX6-pGPD-yeGFP-tURA3-hisG-HO fragment excised from the pTM10-HO_GFP (AB465) plasmid (GenBank: OM069737). These diploids were then sporulated, tetrads were dissected, GFP^+^/NAT^R^ haploid progeny were identified, and their mating types were determined by PCR [41]. These isolates were then propagated on YPD, during which a loop-out of the HO-cassette can occur at a low rate via homologous recombination of the hisG direct repeat. These homologous recombination events leave behind a single copy of the (1130 bp) hisG sequence at the locus, but remove GFP and the NAT drug marker, permitting the future use of these markers. Cells having undergone the desired recombination events were isolated as GFP^-^ cells by FACS, confirmed by PCR using primers AO34 (CTTTGTATCCTTAATAAAATAAAATTCACA) and AO579 (TAAATAGTTACCACAAGGCCATATC) and then tested for nourseothricin-sensitivity.

To facilitate mating and determination of progeny mating type, one of two drug markers (hphMX or natMX)[42] was integrated in the founder strains between *PHO87* and ARS314 (position 197,356 on Chromosome III), which is close to the mating type locus. As a result, in the haploid founder and progeny strains at all levels of the cross, MATa haploids harbored the hygromycin B phosphotransferase (*HPH*) gene conferring hygromycin resistance, while MATα haploids harbored the nourseothricin N-acetyl transferase (*NAT*) gene conferring nourseothricin resistance. These drug resistance phenotypes were used to distinguish diploids and haploids of each mating type. For the final set of eight founder strains, MATa isolates were chosen for four strains (founders 1, 3, 5 and 7) and MATα isolates were chosen for the remaining four (founders 2, 4, 6 and 8).

The mapping population was then derived from the eight founder haploids through three consecutive rounds (“levels”) of pooled mating and sporulation in a “funnel” design (Fig 1B and Fig S1) as follows. For the initial cross, matings were performed in liquid by mixing 150 µl of overnight YPD cultures, each seeded with a single haploid parental colony. The mixtures were incubated at 30°C for 5 hours, without shaking and then plated on YPD+HYG+NAT agar. A single diploid colony was chosen for further crossing. The second and third levels of mating were performed in liquid as above, using two pools of haploid progeny from the previous cross. Specifically, 150 µl of YPD supplemented with the appropriate drug selection was seeded with individual haploid segregants (n=576 strains/mating type) and grown overnight. 10 µl aliquots were then pooled and washed to remove the selective drug. The washed pools of the two haploid populations were mixed and plated directly onto YPD agar. Plates were grown overnight at 30°C, forming lawns which were scraped, washed, and then diluted into YPD+HYG+NAT liquid medium and grown, to select for diploids. For all levels of the cross, heterozygous diploids were grown in YPD+HYG+NAT liquid medium to exponential phase, washed, and then sporulated in Tong and Boone enriched sporulation media [43]. Once sporulated, tetrads were disrupted and individual haploid recombinant progeny were isolated using a FACS-based method previously described.[44] Briefly, 1 ml of the culture was stained with 1 µg/mL DiBAC_4_(5) [bis-(1,3-dibutylbarbituric acid) pentamethine oxonol] (AAT Bioquest 21410), a fluorescent vital dye which selectively stains tetrads and dead cells. To digest the ascus surrounding each tetrad, stained cells were treated with zymolyase 100T and tetrads were disrupted using sterile 0.5 mm diameter glass beads (Biospec Products CAT#11079105) resulting in a population of single spores with very few dyads, triads or intact tetrads. A population of single spores was identified using a Sony LE-SH800 cell sorter configured with a 100 µm microfluidic sorting chip, the 488 nm laser for FSC/BSC detection, and the 561nm laser and 617/30nm bandpass filter set for detecting fluorescence. Individual spores were sorted using the Ultra-Pure setting into single wells of a 96-well plate containing liquid YPD. After growth in YPD, each sorted plate was then replica pinned to both YPD+HYG (to identify MAT**a** cells) and YPD+NAT (to identify MATα cells). Strains that were resistant to both nourseothricin and hygromycin were discarded.

A final set of 5,742 MAT**a** haploid cells and 5,742 MATα haploid cells were collected at the end of the funnel process, for a total mapping population of 11,484 haploid strains. Strains were stored in a 15% glycerol solution at -80°C in 96-well plates, one strain per well.

### Determining ploidy and haplotype of progeny strains

Haplotypes of the 11,484 haploid progeny in the final level of the funnel cross were inferred based on low coverage (median 3.8-fold) WGS as follows. Note that a total of 92 strains failed sequencing quality control, resulting in the final population of 11,392 fully haplotyped strains. Genomic DNA was extracted in 96-well plate format by the method of Drumonde-Neves, et al. [45] and quantitated via a SYBR Green/spectrophotometer assay [46] to obtain a plate-averaged DNA concentration which was then used to dilute the plate concentration to a range optimized for the PlexWell library prep (target = 10 ng template DNA per library prep reaction). Libraries were prepared using the PlexWell 384 Library Preparation Kit (SeqWell Inc., Beverly, MA) according to the instructions in User Guide v20190813. Using this method, each strain was first individually tagged with P7 primers containing a well-specific i7 index using a limited amount of transposase, which serves to normalize the samples. Then, wells of a plate were pooled and tagged using a transposase with a P5 primer containing a plate-specific i5 barcode. Finally, the pool was magnetic-bead-purified and PCR-amplified to enrich P7/P5-tagged fragments and to create the multiplexed library for Illumina sequencing. Paired end (75 x 75 bp, except for 348 strains with 38 x 37 bp) library sequencing was then performed on an Illumina NextSeq 500.

After demultiplexing, sequencing reads underwent adaptor trimming and were aligned to the S288c reference genome as described above for sequencing of the founder strains. A pileup file was then produced for each strain, using the SAMtools (v0.1.19) “mpileup” command with the -C 50 parameter and the BCFtools (v 0.1.17) “view” command with the -c flag (consensus caller). The full set of 11,484 strains was then filtered to remove any with low sequencing coverage (≤12k reads), leaving 11,392 sequenced strains. Mean sequencing depth per chromosome was used to estimate chromosome copy number and to identify aneuploid strains (Table S1).

For each strain in this set, at each position identified in the founder biallelic SNV table, the number of reads matching each of the two alleles was counted from the pileup file for that strain. These counts (Files S2 and S3: https://doi.org/10.5281/zenodo.17362654) were then used to infer the haplotype state of the strain at each of the SNV positions by fitting a hidden Markov model (HMM) to the allele counts using the Viterbi algorithm. The HMM had eight states, corresponding to the eight founder haplotypes. At each SNV site, the emission probabilities for each haplotype were set to 0.99 for the expected allele from that founder strain, with the incorrect allele probability set at 0.01 (error probability). The transition probabilities between haplotype states at each adjacent pair of SNV sites were calculated by combining the expected recombinant fraction per meiosis (R) with the structure of the cross. Specifically, for each pair of adjacent SNVs, R was calculated by first converting the physical distance between the SNVs into genetic distance using a conversion of 3 crossovers per megabase [47]. Then, the genetic distance (m) was converted into R using Haldane’s mapping function, where R = 0.5x(1-e^-2m^). Finally, the structure of the cross was used to determine the transition probabilities (for chromosomes with copy number 1), as a set of equations based on R. For instance, for two adjacent markers in a mapping population strain to both have haplotype 1, no crossover can have occurred in the interval between them in any of the 3 levels of crossing, so this transition probability is (1-R)^3^. Similarly, for a marker to have haplotype 1 and its neighbor marker to have haplotype 2, a crossover must have occurred in the first level of the cross, but no crossover in the remaining two levels, corresponding to R(1-R)^2^. The full set of equations is given in Table S2. The final haplotype inferred for each strain is given in File S4 (https://doi.org/10.5281/zenodo.17362654). A separate set of transition probabilities was used for disomic chromosomes (Table S3).

Pooled sequencing and allele counting of intermediate population G_1_5-6 (Fig 1B) was carried out as described above for individual mapping strains, but at a median sequencing depth of 392-fold at marker positions.

### Scaffolding Marker Subset

A subset of markers was chosen to provide a genome wide “scaffold”, with markers spaced ∼2kb apart so that causative variants should generally be within 1kb of a scaffolding marker. This reduces the number of markers at which statistical tests are carried out while maintaining power and resolution. The set of markers was chosen starting with the first marker on each chromosome, taking the first marker ≥2kb away and repeating until the entire genome was covered. The final scaffold has 5,479 markers (File S5).

### Identifying Mapping Strains with Shared Ancestors in the Intermediate Funnel Cross Populations

To assess bottlenecking, we set out to identify mapping strains descended from the same strain in one of the intermediate funnel cross populations. First, for each pair of mapping strains, regions of the genome descended from the same intermediate cross population were identified as follows. Consider shared ancestry in intermediate population G_1_1-2 (Fig 1B; equivalent to HA1/HA2 in Fig S1), produced by crossing founders 1 and 2. Let x and y be the haplotypes of the two mapping strains being compared. Regions with G_1_1-2 ancestry in both strains are identified as x,y ϵ [23]. For the remaining G_1_ ancestral populations, the test sets used were {3,4}, {5,6} and {7,8}. Regions with shared G_2_ ancestry in both mapping strains were identified using the test sets {1,2,3,4} and {5,6,7,8} (Fig 1B).

For each pair of mapping strains and each intermediate population, haplotype discordance was then measured across the shared subset of the genome. Discordance was calculated as the proportion of markers with non-matching haplotypes in these regions. For aneuploid strains, discordance was calculated only using monosomic chromosomes. Discordance values of 0 are expected when strains share a single ancestor in the intermediate population, otherwise values of 0.5 (G_1_ populations) or 0.75 (G_2_ populations) are expected. To identify groups of strains with a single shared ancestor, hierarchical clustering using the “complete” method was applied to the discordance distances, using the “hclust” command in R.

### Relationship Between Physical and Genetic Distances

For two markers on the same chromosome to have the same haplotype value in a mapping population strain, there must have been no net crossing over between them in any of the three levels of the funnel cross. Therefore, I = (1-R)^3^, where I is the proportion of strains in which the two markers have the same haplotype (identity), and R is the recombinant fraction between those markers per meiosis. In Haldane’s mapping formula, R = 0.5x(1-e^-2m^), where m is the genetic distance between two markers. Therefore, I = (1-0.5x(1-e^-2m^))^3^. By comparing the observed value of I to the value predicted by this formula, a single physical-to-genetic distance conversion coefficient was then estimated by maximum likelihood. Specifically, a range of conversion coefficient values was used to predict I from the physical distance between markers. The value that minimized the residual variance after regressing the observed value of I onto the predicted value was then identified.

### Calculation of LOD Scores in QTL Mapping

For each phenotype, at each tested position, the LOD score was calculated as Log_10_P(B)-Log_10_P(A), where P(A) is the likelihood of the observed data assuming a single normal distribution and a single population mean and P(B) is the likelihood of the observed residuals, assuming normality, after fitting a separate mean for the 8 subpopulations defined by the haplotype at the current marker.

### Single-Marker Deviations from Expected Haplotype Frequencies

Significant deviations from expected haplotype frequencies at the single marker level were identified as follows among the haploid mapping strains. At each marker (except for those on Chromosome IX), the expected proportion of each haplotype in the mapping population is 1/8. Because of the extra copy of Chromosome IX in population G_1_5-6 (Fig 1, 2C, 2D and 2E), expected haplotype frequencies are different for markers on that chromosome. Specifically, the expected frequencies of haplotypes 1, 2, 3, 4, 7, 8 are 1/9 for all Chromosome IX markers, while the expected frequencies of haplotypes 5 and 6 vary between 0, 1/9, 2/9 and 3/9, depending on the region (due to several LOH events on Chromosome IX that occurred along with the increase in chromosome copy number).

We compared the observed haplotype counts at each marker to these expectations using the Chi-square goodness-of-fit test. For its null distribution, this test assumes a Poisson distribution of counts around a mean equal to the expected count (in our case, usually 1/8 of the total number of strains). In the Poisson distribution, the mean is equal to the variance. However, because our final mapping population is much larger than the intermediate cross populations (Fig S1), we anticipated that the count variance in the mapping population would be inflated relative to the mean, producing inflated Chi-square and significance values from the Chi-square test. To test this, we examined the central 95% of ordered counts, which should reflect the null distribution under the assumption that sites under selection are rare. A Q-Q plot of these counts against a simulated Poisson distribution with the same mean confirmed a linear relationship but with a much higher variance in the real counts (S2A Fig). Based on this relationship, we identified the divisor (31) that best transforms the central 95% of ordered counts back to a Poisson distribution (slope closest to 1 for regression against a simulated Poisson distribution) (S2B Fig).

The adjusted counts were used to calculate a p-value for goodness-of-fit at all scaffolding markers. The Holm method was then used to calculate a multiple-hypothesis corrected 0.05 p-value cutoff from these nominal values.

### Pairwise deviations from expected haplotype frequencies

The Chi-square test of independence was used to identify pairs of markers with significant deviation from haplotype independence. Because of the potential for the cross structure to inflate the Chi-square statistic, we examined the bottom 95% of observed inter-chromosomal Chi-square values, which should reflect the null distribution under the assumption that non-independence between (unlinked) markers is rare. A Q-Q plot of these counts against a simulated Chi-square distribution with the same degrees of freedom, confirmed that (null) Chi-square values are indeed elevated in our data (Fig S3). The strong log-linear relationship allowed us to transform the data back to the scale of the expected null distribution. A separate log-linear fit was used for pairwise interactions involving Chromosome IX (Fig S3).

A Chi-square value corresponding to a multiple-hypothesis corrected 0.05 p-value cutoff was then calculated from the transformed inter-chromosomal pair data using the Holm method.

## Results

### Choice of founder strains

To maximize the genetic diversity of our mapping population, we chose eight haploid “founder” strains isolated from diverse geographic and environmental sources (Fig 1A and Table 1), with an emphasis on less extensively studied, non-European isolates. Four of these strains were isolated in China, where *Saccharomyces cerevisiae* originates and where genome diversity is greatest [9, 10, 22]. Three of the Chinese strains were isolated from primordial forests (tropical, subtropical and temperate) and one from a persimmon orchard. Among the non-Chinese strains, one was isolated from soil in the United States, one from soil in South Africa, one from Nigerian palm wine and one from coconut wine in the Philippines (Table 1). WGS of these eight founder strains showed that they capture a large fraction of the currently known genetic diversity in the global population of *S. cerevisiae* (Materials and Methods) [9], including 32% of biallelic single nucleotide variants (SNVs) with a minor allele frequency (MAF) greater than 0.005 and 56% of biallelic SNVs with a global MAF greater than 0.05.

Using the founder WGS data, we identified 276,774 high-quality biallelic SNVs among the founder strains, a density of ∼1 SNVs per 44 bases, with only 0.031% of neighboring SNVs separated by more than 2 kb (File S1: https://doi.org/10.5281/zenodo.17362654). Among these SNVs, 77.4% of minor alleles are unique to a single founder strain, 13.0% are shared by two founders, 6.73% by 3 founders and 2.8% show a 4:4 distribution.

### Mapping population construction

To construct our mapping population, we used an eight-parent funnel cross with three levels of mating (Fig 1B). To initialize the cross, haploid *hoΔ0* founder strains were generated from the original diploid strains. MAT**a** isolates were chosen for four strains (Founders 1, 3, 5 and 7) and MATα isolates were chosen for the remaining four (Founders 2, 4, 6 and 8) (Materials and Methods). We avoided synthetic deletions of metabolic genes (auxotrophic markers) in the founder strains as these can have unintended effects on a range of phenotypes and tend to dominate QTL landscapes, confounding detection of naturally occurring variation [48, 49]. Instead, drug resistance genes were inserted close to the mating type locus, encoding hygromycin resistance in the MAT**a** founders and nourseothricin resistance in the MATα founders. These markers enabled haploid strains of each mating type to be identified at each step in the funnel cross, and allowed diploid strains to be distinguished from haploids, facilitating mating (Materials and Methods).

The mapping population was derived from the eight founder haploids through three consecutive rounds (“levels”) of mating and sporulation as outlined in Fig 1B with details in Fig S1. The second and third rounds of mating were carried out in pools, each consisting of 576 strains of a single mating type derived from spores isolated using a FACS-based approach [44] that enables meiotic progeny to be isolated at scale (Materials and Methods). The large pool sizes prevented bottlenecking at intermediate steps of the funnel, maintaining genetic diversity and avoiding population structure in the final mapping population. At the end of the funnel process 5,742 MAT**a** haploid cells and 5,742 MATα haploid cells were collected. Thus, the entire mapping population (prior to filtering for sequencing coverage) consists of 11,484 haploid strains, each a recombinant mosaic of the eight founder haplotypes. Unlike mice or other obligate diploid organisms, these strains did not require further inbreeding to achieve homozygosity.

### Individual strain ploidy and haplotype

Each of the 11,484 strains in the mapping population underwent low coverage (median 3.8-fold) whole genome sequencing (Materials and Methods). Strains with insufficient coverage (<=12k reads) were then removed, leaving 11,392 successfully sequenced strains in the mapping population. For these strains, we then used relative depth of sequencing coverage per chromosome to estimate copy number (Table S1). This allowed us to identify aneuploid strains (Materials and Methods), which were set aside at this point (see below for discussion of this population), leaving 9,351 euploid (haploid) strains.

Genetically, each strain in the mapping population is a recombinant mosaic of the eight founder haplotypes. We inferred this recombinant haplotype for each euploid mapping strain using the sequencing data as follows. First, the number of reads supporting each of the two possible alleles at the ∼270k biallelic SNVs identified in the founder population were counted (Files S2 and S3: https://doi.org/10.5281/zenodo.17362654). Then, a hidden Markov model was applied to these counts, with eight states representing the eight founder haplotypes. In the model, the alleles present in each of the founder strains were used to set allele emission probabilities for the reads at each SNV. Both estimated linkage between adjacent SNVs and the structure of the funnel cross were used to set transition probabilities between haplotype states (Materials and Methods). Applying this model allowed us to infer the founder haplotype (encoded as values 1:8) of each mapping strain at each SNV position (File S4: https://doi.org/10.5281/zenodo.17362654). The full genotype of each mapping strain can then be derived unambiguously from these haplotypes.

### Assessing the mapping population using a set of ∼5,500 “scaffolding” markers

We next used the haplotyping data to assess three parameters of the euploid mapping population. First, we assessed genome-wide haplotype representation, quantifying how well the genetic diversity of the founder strains is maintained at each genomic position in the mapping population. Second, we assessed linkage between genomic regions, allowing us to identify any genomic rearrangements in the founder strains, relative to the reference (S288c) genome. Finally, we assessed the extent of shared intermediate strain ancestry among the mapping strains, quantifying the degree of population structure. To carry out these analyses, we identified a “scaffolding” subset of 5,479 markers, spaced approximately 2 kb apart, drawn from the full set of ∼270k markers (File S5). This scaffold densely and evenly samples the genome, but greatly reduces the number of markers that need to be assessed statistically, relative to the full set of ∼270k SNVs. Excluding the telomeric regions beyond the first and last marker on each chromosome, 97.7% of the genome is within 2kb of a scaffolding marker.

### Haplotype representation

The proportion of mapping population strains with each founder haplotype was calculated for each of the scaffolding markers. As expected, the mean proportion for each haplotype genome-wide closely approximated 12.5% (1/8) (Fig 2A), with standard deviations ranging from 1.7% (haplotype 2) to 3.0% (haplotype 6). For every haplotype, representation at a level greater than 5% was seen across >95% of the genome (Fig 2B).

**Fig 2.**
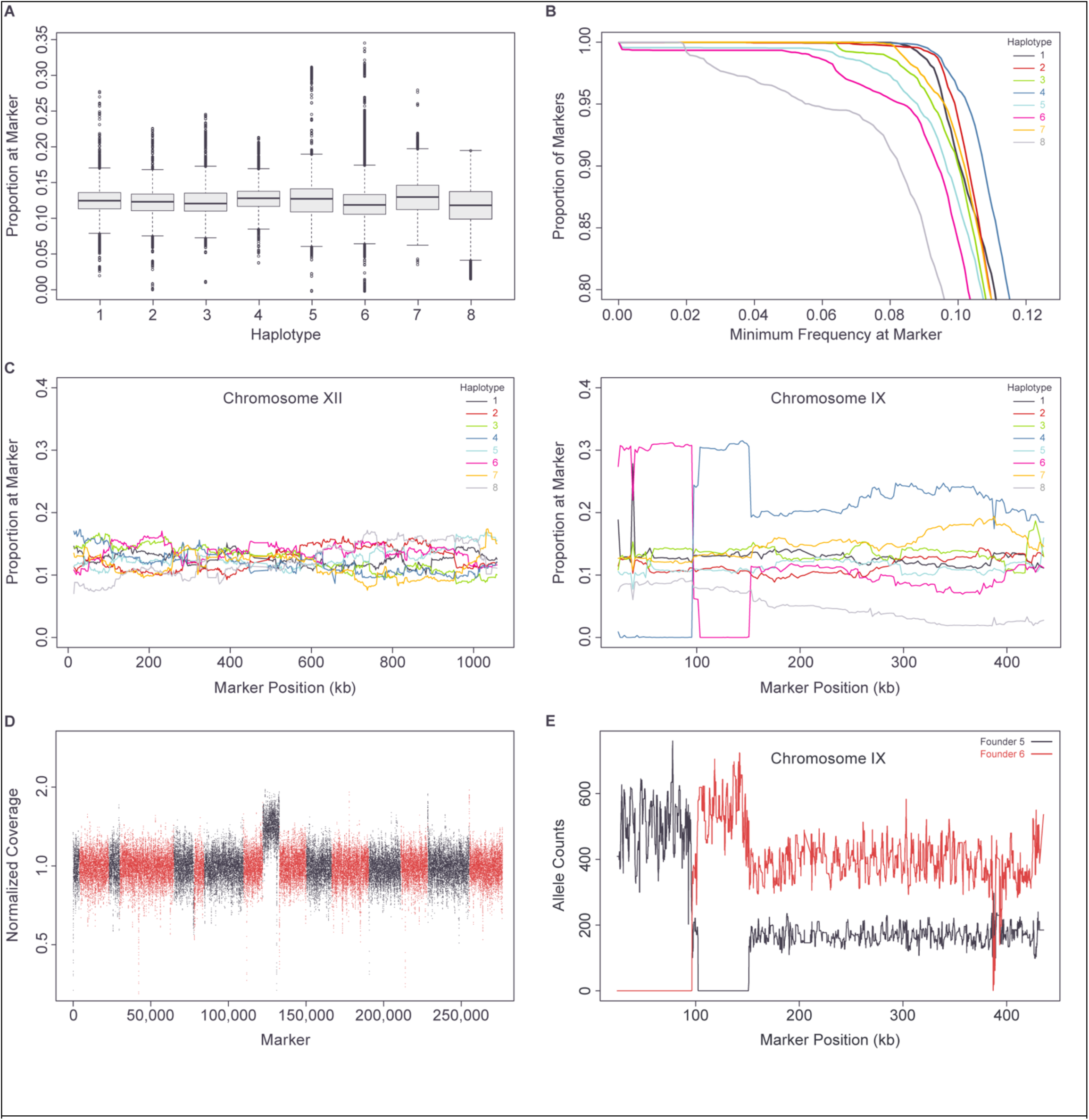
Genome-wide haplotype representation. (A) Boxplot of haplotype proportion at each of the ∼5.5k scaffolding markers. (B) Proportion of scaffolding markers with representation of each haplotype above cutoff. (C) Haplotype proportion by scaffolding marker position across Chromosomes XII and IX. (D) Genome-wide sequencing depth in haploid population G_1_5-6 measured in 31-bp sliding windows, normalized by genome median. Chromosomes indicated in alternating red and black. (E) Allele counts on Chromosome IX, at markers polymorphic between founders 5 and 6, from genome-wide sequencing of haploid population G_1_5-6.

Haplotype frequencies across a typical chromosome (Chromosome XII) are shown in Fig 2C. In contrast, an unusual distribution of haplotype frequencies was seen across Chromosome IX (Fig 2C), with an ∼50% elevated frequency of haplotypes 5 or 6 relative to other haplotypes, one region with complete loss of haplotype 5 and another region with complete loss of haplotype 6. Pooled sequencing of haploid population G_1_5-6, formed by crossing founders 5 and 6 (Fig 1B) identified a 50% increase in the copy number of Chromosome IX relative to the rest of the genome (Fig 2D; Materials and Methods). In addition, in this pooled sequencing, a 2:1 ratio of haplotype 6 to haplotype 5 was seen across most of the chromosome, along with two regions showing loss of either haplotype 5 or haplotype 6 representation (Fig 2E), matching the pattern in the final mapping population (Fig 2C). Haploid founder strains 5 and 6 are euploid, therefore our data are consistent with a duplication of the haplotype 6 copy of Chromosome IX in the diploid formed by mating them together (“DA3” in Fig S1), i.e. trisomy IX, along with two LOH events on the left arm of Chromosome IX. This extra copy of Chromosome IX at the top of the funnel cross, made up of haplotypes 5 and 6, explains the elevated frequency of these Chromosome IX haplotypes in the final euploid mapping strains (Fig 2C), as well as the high frequency of disomy IX among the aneuploid strains (see discussion of the aneuploid strains, below).

### Linkage behavior

To estimate genetic linkage between genomic regions in the euploid population, for each of the scaffolding markers, the recombinant haplotype fraction (proportion of non-matching haplotypes) was calculated against all of the remaining markers. The resulting data (Fig 3A) show little evidence of linkage patterns other than those expected by physical proximity between markers on the reference genome. That is, there is little sign of genome rearrangements in the founder strains relative to the S288c reference. The only exceptions involve a small number of telomeric markers.

**Fig 3.**
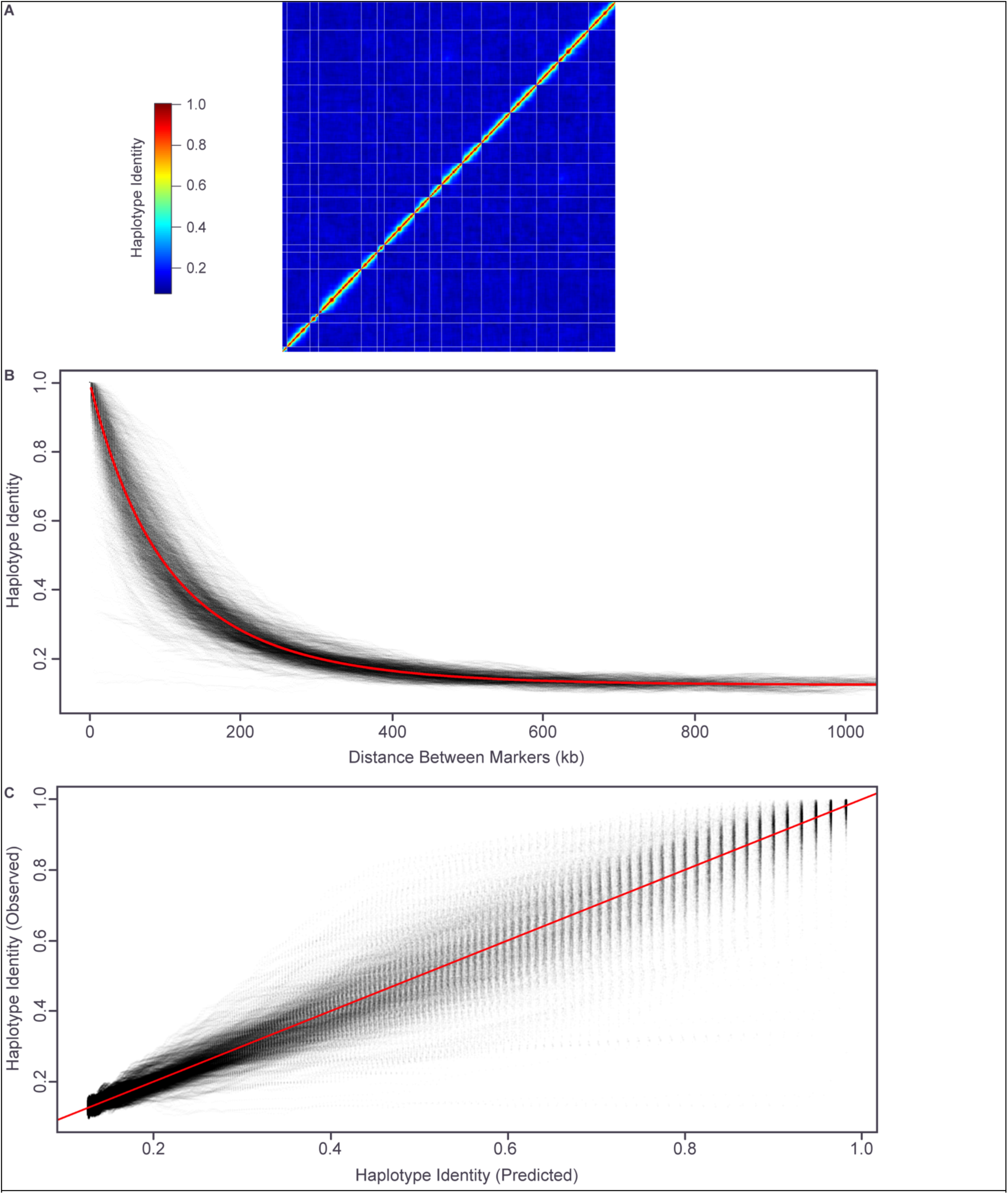
Genome-wide linkage behavior. (A) Genetic linkage as proportion haplotype identity between each pair of scaffolding markers ordered from beginning of Chromosome I to end of Chromosome XVI. White lines separate chromosomes. (B) Observed (grey) and fitted (8.7 crossovers per megabase, red) relationship between physical distance and linkage for each pair of scaffolding markers on the same chromosome. (C) Observed linkage versus fitted linkage (based on 8.7 crossovers per megabase) for each pair of scaffolding markers on the same chromosome.

We also modelled the relationship between physical and genetic distances, assuming a Poisson distribution of crossovers (no interference) and a linear relationship between physical distance and the mean number of crossovers (genetic distance) (Materials and Methods). A genome-wide average value of 8.7 crossovers per megabase (after 3 rounds of meiosis) provided the best fit (Fig 3B and 3C).

### Population structure

Population structure has the potential to generate spurious genotype-phenotype correlations. Our funnel cross was designed to minimize such structure in our mapping resource by using large pools of strains in the intermediate crossing steps, avoiding bottlenecking. However, processes such as variations in mating efficiency or growth could still lead to some degree of bottlenecking, with specific intermediate cross strains, and their associated haplotypes, contributing disproportionately to the final mapping population. Therefore, to quantify population structure and bottlenecking, for each pair of strains in the final (euploid) mapping population, we determined if they shared an ancestral strain in any of the intermediate haploid populations involved in the funnel cross.

Our approach is based on identifying regions of shared intermediate population ancestry in each pair of strains. For each strain in the final mapping population, the haplotype of each marker identifies the inheritance path of that region through the funnel cross structure. For example, any marker with haplotype 1 was inherited from founder 1, via intermediate haploid populations G_1_1-2 and G_2_1-4 (Fig 1B). Similarly, each intermediate haploid population is responsible for inheritance of a specific set of founder haplotypes in the final progeny strains. For example, in every progeny strain, all markers with haplotypes 1-4 were inherited via intermediate population G_2_1-4 (Fig 1B) and all markers with haplotypes 1-2 were inherited via intermediate population G_1_1-2 (Fig 1B). In each progeny strain this allows us to identify “G_1_1-2 inherited”, “G_2_1-4 inherited” regions etc.

Having identified regions in every strain descended from each intermediate population (e.g. population G_1_1-2), for each pair of strains we can then identify the common regions derived from that population in both strains. If the two strains share a single ancestor in the intermediate population of interest, their haplotypes will be identical across the shared region. Otherwise, for unrelated strains, the proportion of identical haplotypes in the shared region should be 0.5 (for intermediate “G_1_” populations consisting of two haplotypes, Fig 1B) or 0.25 (for intermediate “G_2_” populations consisting of four haplotypes, Fig 1B). For each pair of strains, and for each intermediate population, we quantified this as a discordance “distance” (proportion non-matching haplotypes in the shared region, Materials and Methods).

Clustering the results for each of the intermediate haploid populations (Materials and Methods) reveals only a few small clusters with shared ancestry and therefore, little evidence of bottlenecking at the level of the “G_1_” or “G_2_” intermediate populations (Fig 4).

**Fig 4.**
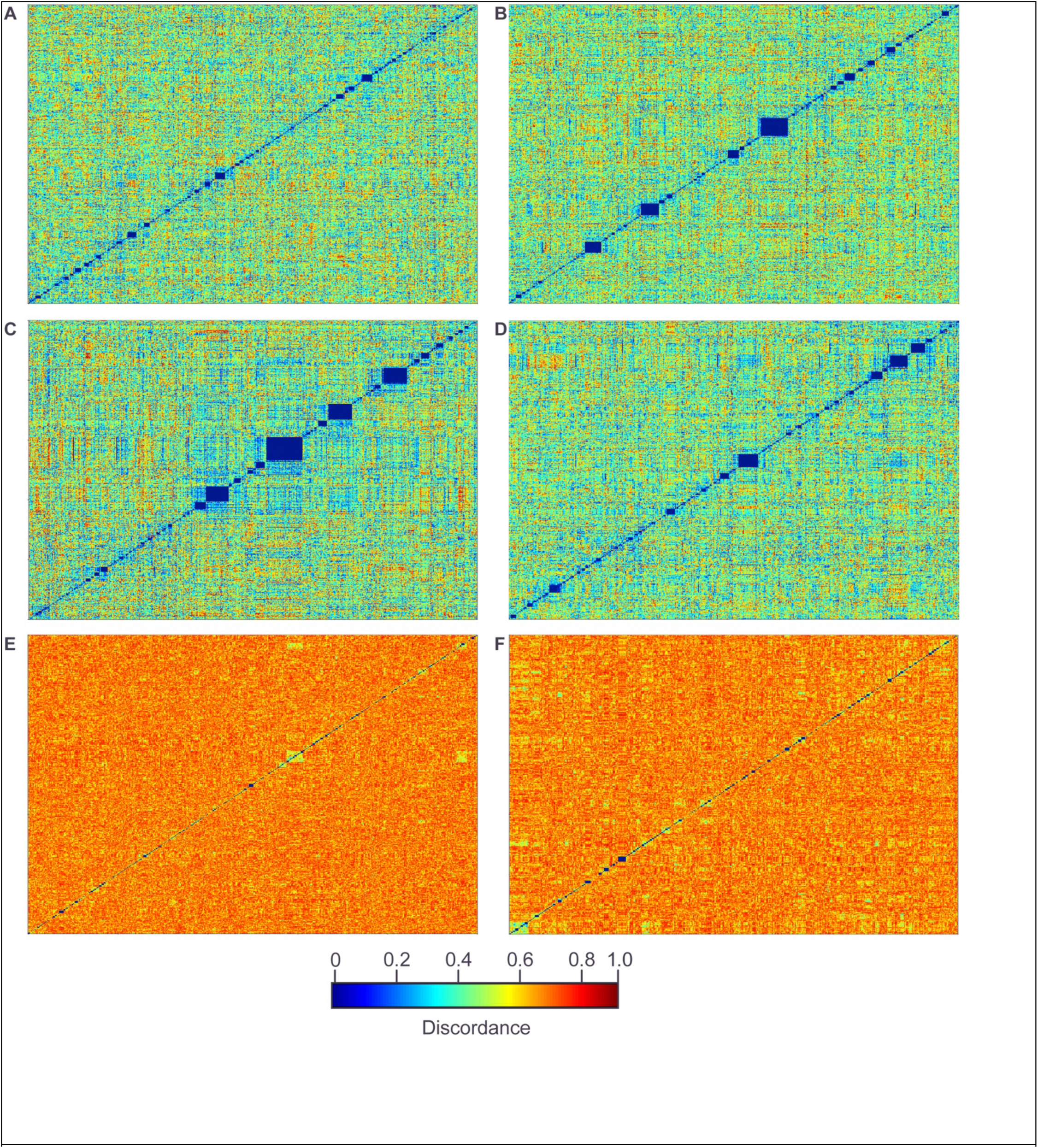
Quantifying shared ancestry in intermediate cross populations. All progeny strains clustered by pairwise haplotype discordance relative to each intermediate population. Shared ancestry is expected to produce a discordance value of 0. (A) Population G_1_1-2. (B) Population G_1_3-4. (C) Population G_1_5-6. (D) Population G_1_7-8. (E) Population G_2_1-4. (F) Population G_2_5-8. The discordance scale applies to all plots in the figure.

These results both quantify the ancestral relationship between strains in the mapping population and demonstrate that the degree of population structure in our mapping population is very low.

### Scaffolding markers as an efficient and highly powered resource for linkage mapping

The ∼5.5k set of scaffolding markers was chosen to cover the yeast genome evenly and densely, and such that 97.7% of the genome (excluding telomeric regions) is within 2kb of a scaffolding marker. Therefore, at least one scaffolding marker should be closely linked genetically to any causative locus. Because of these features, the scaffolding marker set should behave as an efficient, but still highly powered, alternative to QTL mapping with the full set of ∼270k high-quality SNVs. To compare the mapping power of the scaffolding set to that of the full set of markers, we first randomly chose 1,000 out of the full set of ∼270k markers to be QTL. We then generated simulated phenotypes for all of these causative markers across a range of heritability values (*h*^2^ > 0,), i.e. varying the proportion of the total phenotypic variance explained by the marker genotypes. Each simulated phenotype had a mean of 0 and a variance of 1 and residuals were normally distributed. For each simulated phenotype, we then calculated the LOD at the causative locus and at the scaffold marker most closely genetically linked to the causative locus. LOD scores were calculated by regressing phenotype on haplotype (a factor with eight levels) at each marker (Materials and Methods). Our results demonstrate that the linked scaffolding markers show very little reduction in power relative to the causative loci, with a regression estimate of LOD_scaff_ = LOD_caus_^0.9949^ (Fig 5A).

**Fig 5.**
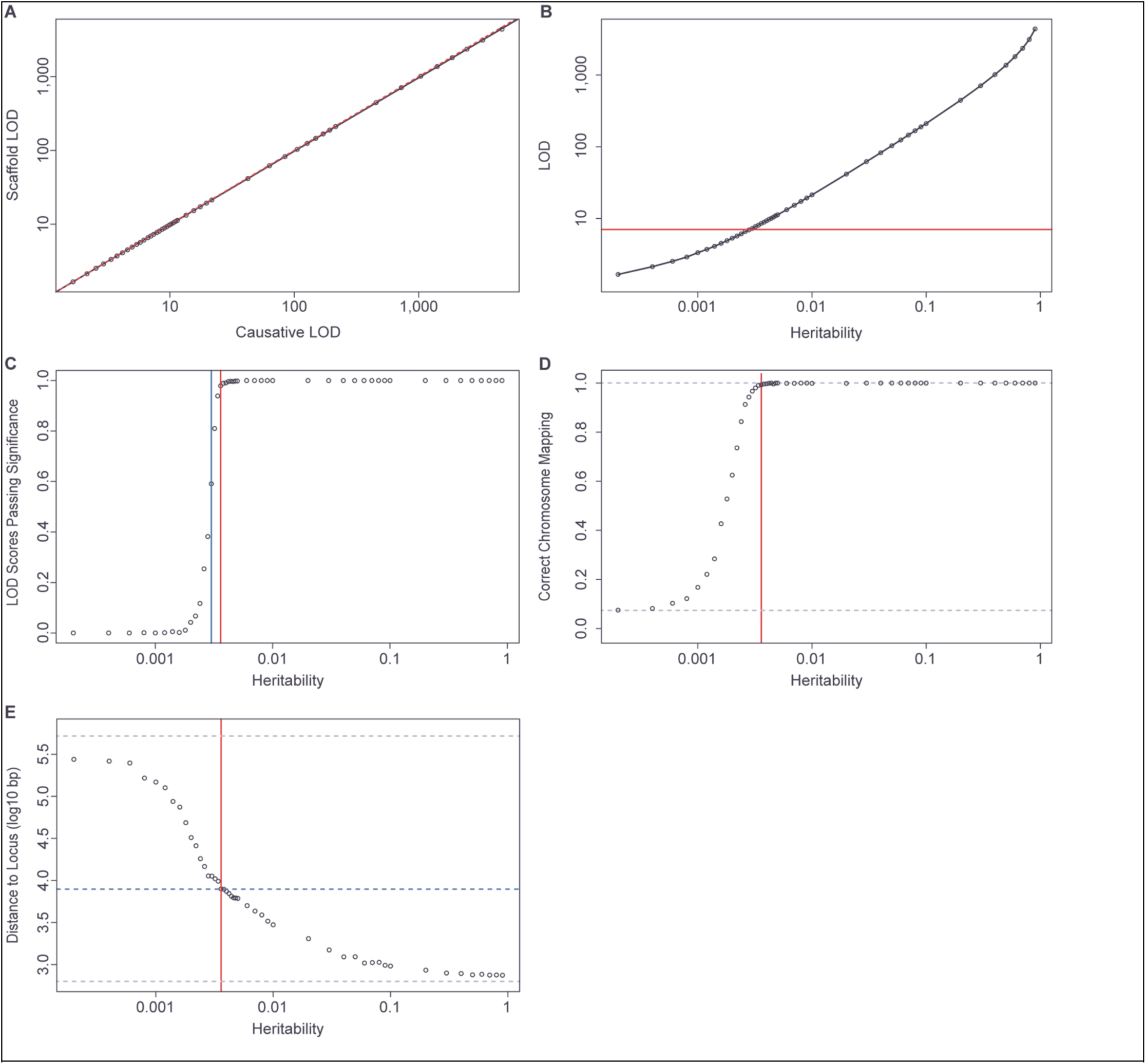
Simulated QTL mapping power and resolution. (A) Mean LOD scores of causative markers versus most closely linked scaffold markers. Regression line in black, dashed 1:1 line in red. (B) Causative marker heritability versus mean LOD at the most closely linked scaffolding marker. Red line indicates the 5% type I family-wise LOD error threshold. (C) Relationship between heritability at the causative marker and the proportion of LOD scores exceeding the 5% significance threshold at the most closely linked scaffolding marker. Blue and red lines, respectively, identify the heritabilities (0.3% and 0.36%) at which >=50% and >=95% of scaffolding markers exceed the significance threshold. (D) Proportion of highest-scoring scaffolding markers mapped to same chromosome as causative maker, as a function of causative marker heritability. Lower grey dotted line indicates the proportion expected by chance (minimum possible) and the upper dotted line lies at 1 (maximum possible). Red line represents heritability of 0.36%. (E) Mean distance between causative locus and highest scoring scaffolding marker when on the same chromosome for different levels of heritability. Lower grey dotted line indicates the best possible resolution (distance from causative marker to most closely linked scaffold marker) and the upper dotted line indicates the worst possible resolution (mean distance between causative marker and a random marker on the same chromosome). Red line represents heritability of 0.36% and blue dotted line represents mean distance between the maximal scaffolding marker and the causative marker for that heritability.

### Predicted power and mapping resolution

Based on the similarity in power between the full set of markers and the scaffolding subset, we used the scaffolding markers to assess the power of our mapping population to detect QTL. To this end, we first calculated an empirical 5% type I family-wise LOD error threshold for the scaffolding markers. Specifically, we simulated 1,000 replicates where, for each replicate, we assigned phenotypes to each of the 11,392 segregants under the null hypothesis *h*^2^ = 0 (i.e., the phenotype was sampled from a normal distribution with mean 0 and variance of 1), calculated the maximum LOD score across the ∼5.5k scaffolding markers, and determined the 95^th^ percentile of these maximum LOD score values.

For each of the previously described phenotypes that were simulated with varying levels of *h*^2^ > 0, we then carried out regression across the full set of scaffolding markers and identified the maximum genome-wide LOD score. We refer to the marker with the highest LOD score as the “candidate marker.” The mean candidate marker LOD score achieved significance (at the empirical 5% level) for simulated phenotypes with a *h*^2^ ≥ 0.3% (Fig 5B). The median candidate marker score also achieved significance at this level of heritability, while at heritabilities of 0.36% and above, >95% of individual candidate marker LOD scores achieved genome-wide significance (Fig 5C). Therefore, our results indicate that the scaffolding markers are highly powered to detect QTL with ℎ^2^ ≥ 0.36%.

Comparing the position of the causative loci to that of the corresponding candidate markers allowed us to assess the ability of the scaffolding markers to resolve QTL positions. Consistent with the empirical significance cutoff, the frequency with which the candidate marker lies on the same chromosome as the causative locus declines rapidly at heritabilities below 0.0030 (Fig 5D). At our more highly powered heritability cutoff of 0.0036, the candidate marker is on the correct chromosome 99.3% of the time, with a mean distance to the causative locus of 7.9kb (Fig 5E). This distance falls to <5kb, <2kb and <1kb at heritabilities greater than or equal to 0.007, 0.03 and 0.09, respectively (Fig 5E).

Together, these results demonstrate that mapping with the scaffolding markers is highly powered to detect QTL with heritabilities >0.0036 and to identify the location of causative loci at a resolution of a few kb or less.

### Loci with single and pairwise deviations from expected haplotype frequencies

The repeated steps of mating, sporulation and growth that make up the funnel cross process have the potential to act as a selective regime, enriching for high-fitness alleles and their associated haplotypes and depleting low-fitness alleles and haplotypes. These processes would produce deviations from the background distribution of haplotype frequencies at the single-marker level. Similarly, pairwise interactions between loci could also affect fitness, including through synthetic inviability. Such pairwise interactions would generate statistical non-independence between haplotypes at the two interacting loci. We used the scaffolding markers to identify loci where haplotype frequencies deviate significantly from the null expectation for single markers (generally 1/8 proportion of each haplotype, Materials and Methods). In addition, we also looked for significant levels of haplotype non-independence between all pairs of scaffolding markers (Materials and Methods). These analyses identified a strong signal at the single-marker level on Chromosome IV (Fig 6A) as well as three sets of strong pairwise interactions (Fig 6B), which also had marginal effects detectable at the single-marker level (Fig 6A).

**Fig 6.**
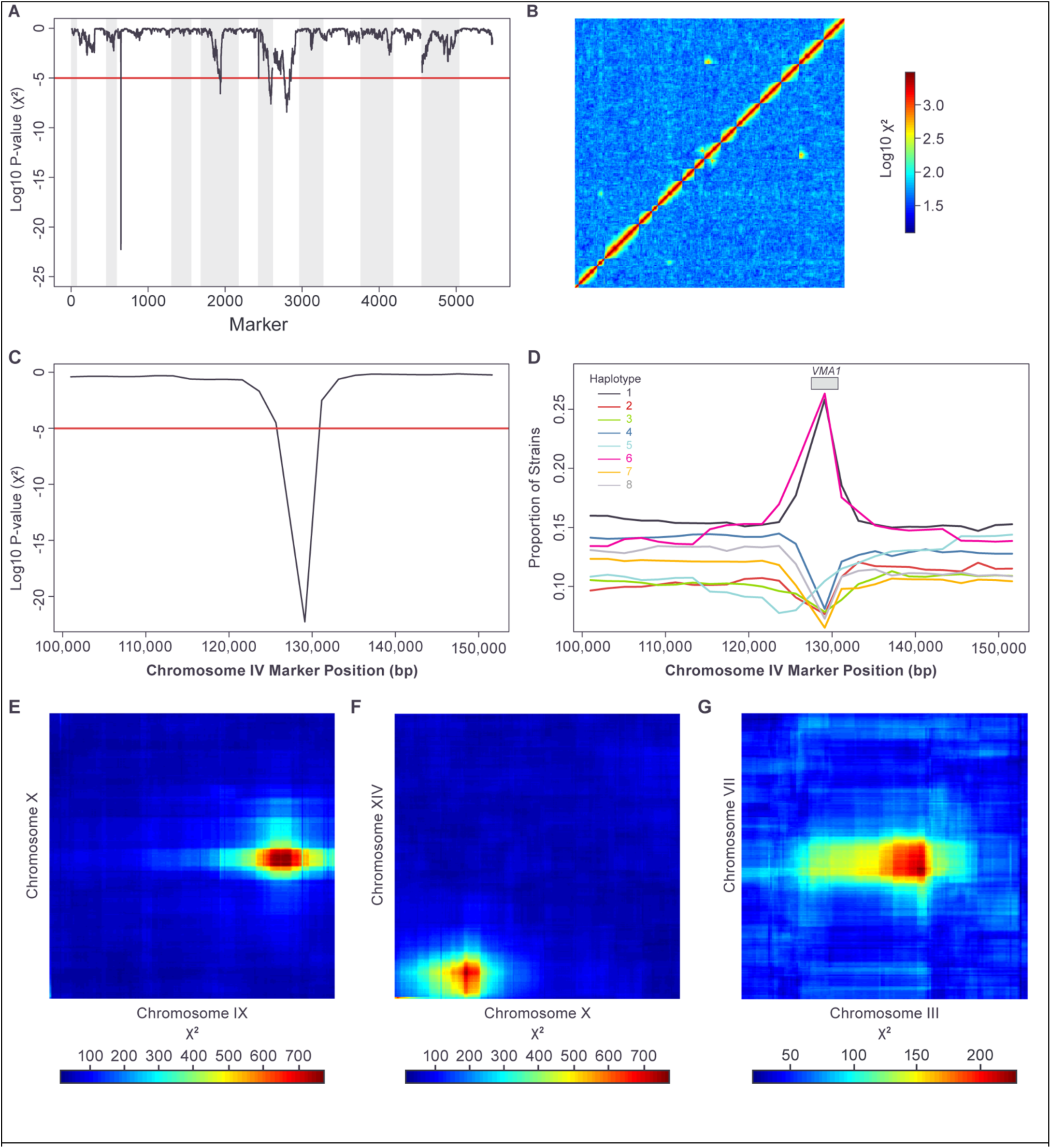
Single-marker and pairwise deviations from expected haplotype frequencies. (A) Nominal Chi-square P-value for single marker deviation from expected haplotype proportions. Multiple-hypothesis corrected 5% significance cutoff (Holm method) shown in red. Chromosomes indicated with alternating grey and white background. (B) Log_10_ Chi-square statistic values for pairwise independence between markers genome-wide. Multiple-hypothesis corrected 5% significance cutoff (Holm method) corresponds to a value of 131 (log_10_ value = 2.12). (C) Closeup of panel a) for region on Chromosome IV around VMA1 gene. (D) haplotype proportions for region on Chromosome IV around VMA1 gene. (E-G) Chi-square statistic values for pairwise independence between markers on the two chromosomes. Multiple-hypothesis corrected 5% significance cutoff (Holm method) is equal to 131.

The single-marker signal on Chromosome IV (Fig 6A) localizes to the *VMA1* gene (Fig 6C) and is caused by a spike in the frequency of haplotypes 1 and 6 over a short region centered on the gene (Fig 6D). This pattern is the expected result of a known gene conversion process driven by a homing endonuclease. Specifically, the *VMA1* sequences of founder strains 1 and 6 encode the site-specific homing endonuclease PI-SceI, while the other six founders have VMA1 sequences that lack the endonuclease sequence (Fig S4). PI-SceI is released from Vma1 by protein splicing and, during meiosis, PI-SceI cleaves *VMA1* sequences that lack the endonuclease sequence, initiating conversion to the endonuclease-containing form via recombination [50]. Our data are consistent with this process occurring in the intermediate funnel cross diploids (DA1, DA3, DB1-4, DC1-2; Fig S1) where an allele of *VMA1* containing the endonuclease (from founders 1 or 6) is present with an endonuclease-free *VMA1* gene derived from any of the other founders.

Three pairs of loci displayed strong and significant deviations from haplotype independence (Materials and Methods). For the first of these pairs, the maximal deviation was for scaffolding markers at positions 361,254 on Chromosome IX (in the region of the *AIM1* gene) and 357,553 on Chromosome X (in the region of the *SDH1* gene) (Fig 6E). Examination of the pairwise haplotype count table revealed a “synthetic viability” pattern for haplotype 8, where there are almost no strains with haplotype 8 at either locus except when both loci have haplotype 8 (Table 2). This pattern suggests inviability when haplotype 8 at either locus is combined with any other haplotype at the other locus. The second pairwise interaction was maximal between marker pairs at positions 204,464 on Chromosome X (in the region of the *MDV1* gene) and 116,372 on Chromosome XV (in the region of the *NDJ1* gene) (Fig 6F) and showed the same kind of “synthetic viability” pattern seen with the first pair of loci, but this time involving haplotype 6 (Table 3). The final pairwise interaction was maximal for markers at positions 201,158 on Chromosome III (adjacent to the mating type locus) and 481,269 on Chromosome VII (in the region of the *PMA1* gene) (Fig 6G). This pairwise interaction displayed a more complex pattern than that seen with the two “synthetic viability” interactions (Table 4).

**Table 2.** Pairwise haplotype strain counts for markers at position 361254 on Chromosome IX and 357553 on Chromosome X.

|  |  | ChrX |  |  |  |  |  |  |  |
| --- | --- | --- | --- | --- | --- | --- | --- | --- | --- |
|  | Haplotype | 1 | 2 | 3 | 4 | 5 | 6 | 7 | 8 |
| ChrIX | 1 | 147 | 160 | 107 | 136 | 162 | 138 | 211 | 0 |
|  | 2 | 187 | 162 | 141 | 172 | 170 | 138 | 235 | 0 |
|  | 3 | 135 | 185 | 253 | 75 | 176 | 186 | 267 | 0 |
|  | 4 | 218 | 146 | 66 | 108 | 158 | 124 | 191 | 0 |
|  | 5 | 106 | 92 | 90 | 79 | 93 | 88 | 128 | 0 |
|  | 6 | 301 | 332 | 274 | 235 | 367 | 270 | 469 | 1 |
|  | 7 | 231 | 226 | 200 | 193 | 215 | 260 | 365 | 0 |
|  | 8 | 1 | 0 | 0 | 1 | 0 | 0 | 0 | 180 |

**Table 3.**
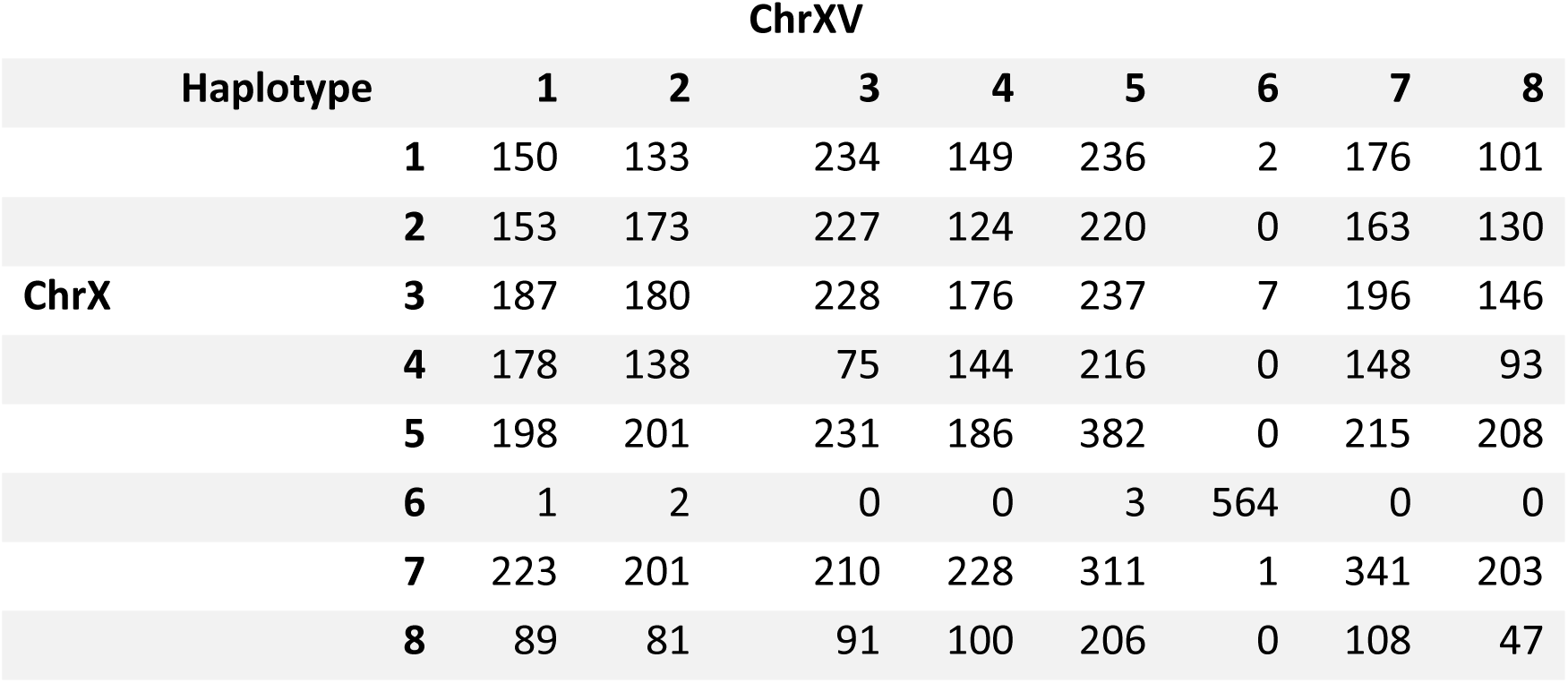
Pairwise haplotype strain counts for markers at position 204464 on Chromosome X and 116372 on Chromosome XV.

**Table 4.** Pairwise haplotype strain counts for markers at position 201158 on Chromosome III and 481269 on Chromosome VII.

|  |  | ChrX |  |  |  |  |  |  |  |
| --- | --- | --- | --- | --- | --- | --- | --- | --- | --- |
| Haplotype |  | 1 | 2 | 3 | 4 | 5 | 6 | 7 | 8 |
| ChrIII | 1 | 88 | 48 | 139 | 374 | 109 | 189 | 166 | 182 |
|  | 2 | 188 | 181 | 61 | 100 | 81 | 64 | 159 | 175 |
|  | 3 | 238 | 210 | 31 | 92 | 87 | 178 | 138 | 146 |
|  | 4 | 110 | 81 | 90 | 335 | 95 | 87 | 240 | 161 |
|  | 5 | 187 | 175 | 95 | 285 | 54 | 13 | 349 | 336 |
|  | 6 | 134 | 107 | 69 | 213 | 97 | 307 | 61 | 52 |
|  | 7 | 95 | 100 | 51 | 160 | 142 | 119 | 78 | 29 |
|  | 8 | 198 | 139 | 84 | 290 | 82 | 54 | 252 | 321 |

Depending on the pattern of interaction, pairwise deviations from haplotype independence can, but do not always, produce marginal haplotype effects detectable at the single marker level. The three pairwise interactions described here colocalize with the significant deviations from expected haplotype frequencies at the single-marker level on Chromosome VII, IX and X (Fig 6A and 6B).

### Aneuploid strains

Finally, we analyzed the aneuploid subpopulation of mapping strains that were separated from the euploid strains earlier in our analysis. Aneuploidy occurs frequently in wild strains and is often well tolerated,[9, 51, 52] and aneuploid strains were quite common among the progeny of the funnel cross, consisting of 2,041 strains (17.9% of the total mapping population, after removing strains with low sequencing coverage). Relative chromosome coverage indicated which chromosomes were disomic in each strain, i.e. no chromosome copy numbers higher than 2 were observed (Table S1). Haplotypes were inferred for monosomic chromosomes as described for the euploid subpopulation, while a separate model was used to infer the haplotypes of disomic chromosomes (Materials and Methods). The final haplotypes for each aneuploid strain are given in File S7 (https://doi.org/10.5281/zenodo.17362654).

Among the 2,041 aneuploid strains, 1,824 (89.2%) displayed patterns of sequence-coverage consistent with disomy of a single chromosome and 206 (10.1%) had disomy of 2 chromosomes (Table S1). Only 15 (0.7%) of strains had disomy of 3 or more chromosomes. Although disomies of all chromosomes were observed in the aneuploid population (Table 5), the most common were of Chromosomes IX (49.7%), X (16.5%) and I (14.5%). Chromosome I is the smallest yeast chromosome and frequently displays copy number changes in natural yeast isolates,[9, 52] implying that is prone to missegregation during cell division. Disomy I is also well tolerated,[52, 53] likely because it perturbs the copy number of only a small proportion of the genome. The high observed frequency of disomy IX is expected given the extra copy of this chromosome inferred to have been present in strain DA3 (Fig 2C, 2D, 2E and Fig S1). Like Chromosome I, disomy of Chromosome IX is also frequently observed in strains not adapted to laboratory conditions.[9, 11, 52] Disomy X co-occurs much more frequently with disomy IX than expected by chance based on their individual frequencies in the full (∼11k) mapping population (Chi-squared test of independence: p<1x10^-16^), with 30.7% of strains disomic for Chromosome X also disomic for Chromosome IX (Table S1). The known mechanism giving rise to disomy IX at the beginning of the funnel cross suggests that disomy X arose preferentially from strains that were already disomic for Chromosome IX. This could occur in one of two ways. First, disomy X could have arisen in a small number of ancestral strains disomic for Chromosome IX, followed by expansion (selective or stochastic) of these lineages. Second, disomy X could have arisen independently multiple times in strains disomic for Chromosome IX, i.e. at an elevated rate in such strains. The first possibility would be detectable as strong bottlenecking in population G_1_5-6 (Fig 1B) among the Chromosome X disomes, which is not observed (Fig S5). Therefore, it appears that disomy X arose multiple times independently, but strongly associated with existing Chromosome IX disomy, suggesting that increasing the copy number of Chromosome X may be adaptive in strains with an extra copy of Chromosome IX.

**Table 5.**
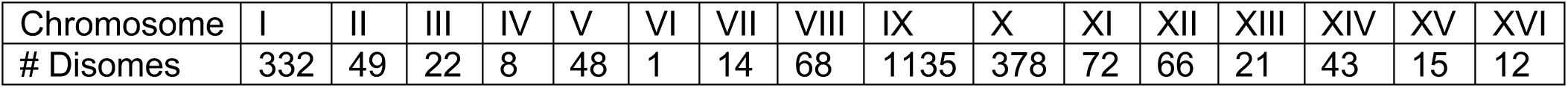
Count of strains disomic for each chromosome.

## Discussion

CYClones, a fully genotyped library of 11,392 recombinant progeny, is presented here as a resource for the yeast community. The strain collection was produced using an eight-parent funnel cross design, starting with fully sequenced haploid founder strains chosen to capture a large proportion of the total yeast population diversity (56% of all biallelic SNVs with a global MAF >5%) [9]. At each iteration of the funnel cross, haploid segregants were isolated using a FACS-based method that enables individual haploid segregants to be sorted by flow cytometry without incorporating a fluorescent tag [44]. As a result, one of the more commonly used drug markers has been preserved, enabling CYClones to be crossed to popular community resources such as the yeast deletion collection [54].

Low coverage sequencing of the CYClones strains was used in conjunction with the known alleles of the eight founder strains at all SNV positions to create a haplotype representation of each haploid segregant at each of the ∼270k SNV positions (∼1 SNV per 44 bases). These haplotypes can either be used directly for purposes such as QTL mapping or can be converted unambiguously into genotypes using the known founder genome sequences. We also present a reduced “scaffolding” subset of ∼5500 markers (spaced ∼2kb apart) that can be used for linkage mapping. This scaffold samples the genome densely and evenly, but greatly reduces the number of markers that need to be assessed statistically, relative to the full set of ∼270k SNVs. Using the CYClones scaffolding haplotypes, we examined linkage behavior, population structure and haplotype diversity in the mapping population. The funnel cross design involved three rounds of meiosis, resulting in approximately 9 crossovers per megabase in each of the CYClones strains. Linkage was consistent with the pattern expected from the reference genome, indicating no large-scale genome rearrangements. The CYClones library exhibits very low levels of population structure, with little evidence of bottlenecking in any of the intermediate cross populations. Examination of haplotype frequencies demonstrated that haplotype diversity was maintained genome-wide, with representation at a level greater than 5% seen across >95% of the genome for all founder haplotypes. In addition, a small number of loci showing significant deviations from expected allele frequencies have been identified, including several pairwise deviations indicative of “synthetic viable” genetic interactions.

CYClones was designed primarily as a powerful tool for QTL mapping, with the large size of the genotyped population and high density of markers providing statistical power to detect loci of small effect and gene-gene interactions. In addition, the density of recombination events associated with three rounds of crossing during strain construction should allow the positions of QTL to be mapped with high resolution. Furthermore, the ∼5500 “scaffolding” markers represent an efficient way of harnessing this power relative to the full set of ∼270k markers, with 97.7% of the genome (excluding telomeric regions) lying within 2kb of a scaffolding marker. Using simulated phenotypes affected by a single genetic locus, we demonstrate that the most closely linked scaffolding marker is almost equally powered to detect a QTL compared to the causative marker itself. Further simulations indicate that the scaffolding markers are highly powered to detect QTL with *h*^2^ ≥ 0.36% and to identify the location of such causative loci at a resolution of a few kb or less. Once a QTL has been detected using the scaffolding markers, its position can be further resolved using local markers drawn from the full set of ∼270k.

We identified a substantial subset of aneuploid CYClones strains containing a variety of disomic chromosomes. These aneuploid strains are fully haplotyped, including the disomic chromosomes. Unlike mammals, where an increase in copy number of most autosomes during meiosis is a fatal event, in yeast aneuploidy is commonly observed in strains isolated from the environment, agricultural products and immunocompromised individuals [5, 6, 8, 9, 11, 22, 52, 55]. Copy number changes in yeast can be an advantageous means of rapidly adapting to environmental stresses, undergoing phenotype switching, or balancing the effect of genetic incompatibilities [56–58]. Like their euploid counterparts, the aneuploid subset of CYClones encompasses a large portion of the genetic variation known to exist in *S. cerevisiae*. Therefore, these segregants represent a singular resource for exploring interactions between copy number and allelic effects on phenotype. While the aneuploid population includes at least one strain (and generally more than a dozen strains) for each of the sixteen possible disomies, a duplication event at the top level of the cross has resulted in >1,000 strains containing a Chromosome IX disomy, making this subset of aneuploids highly powered for characterizing QTL in the context of a specific disomic chromosome.

Because of its large size, genetic diversity, and random assortment of alleles, CYClones will provide a powerful resource for QTL mapping and a range of other purposes. For example, given the density of variant sites in the population and the multiple rounds of meiosis, our library provides a rich dataset for understanding the molecular mechanisms of recombination and chromosomal segregation. CYClones could also be used to explore the genome dynamics of non-reciprocal events associated with retrotransposition (Ty elements) or homing endonucleases, such as the effect of PI-SceI at the *VMA1* locus described in this study. CYClones is also a powerful resource for identifying yeast strains with extreme phenotypes, with utility for industrial, pharmaceutical, and food science applications. In short, the CYClones collection is a powerful and flexible tool for the research community.

## Acknowledgements

We thank Drs. Feng-Yan Bai and Justin Fay for providing yeast strains. We thank Barak Cohen, Justin Fay, and Robi Mitra for helpful discussions. This work was funded by an NIH/NIGMS award (R01 GM117119) to A.M.D. and J.M.A., award R01 GM110068 to J.M.A., and institutional funding from the Pacific Northwest Research Institute to A.M.D.

## Author Contributions

Experiments were designed by A.M.D., J.M.A., G.A.C., R.S.L., and A.E.C. Experiments were performed by R.S.L., T.S.M., M.S.T. and J.A. under the supervision of A.M.D. Analyses were conducted by G.A.C., and A.E.C. under the supervision of A.M.D. and J.M.A. Software was written by G.C. The manuscript was written by G.A.C., R.S.L., A.S. and A.M.D. and incorporates comments by all other authors.

## Data availability

Sequence reads for the eight parental founder strains are deposited in the NCBI Short Read Archive (accession PRJEB80495), and sequence reads for the progeny strains are available under accession PRJEB72283. All supporting datasets are publicly available as supplementary files on Zenodo at https://doi.org/10.5281/zenodo.17362654 (CC BY 4.0 license): parent marker genotypes (File S1), raw marker read counts (Files S2 and S3), inferred euploid and aneuploid haplotypes (Files S4 and S7), the 2 kb marker scaffold (File S5) and an R script (File S6) that reproduces all analyses described in the Results, starting from the haplotype table. Full details of haplotype inference, including emission and transition probabilities and use of the Viterbi algorithm, are described in the Materials and Methods to enable independent reimplementation if desired.

---

**Fig S1.**
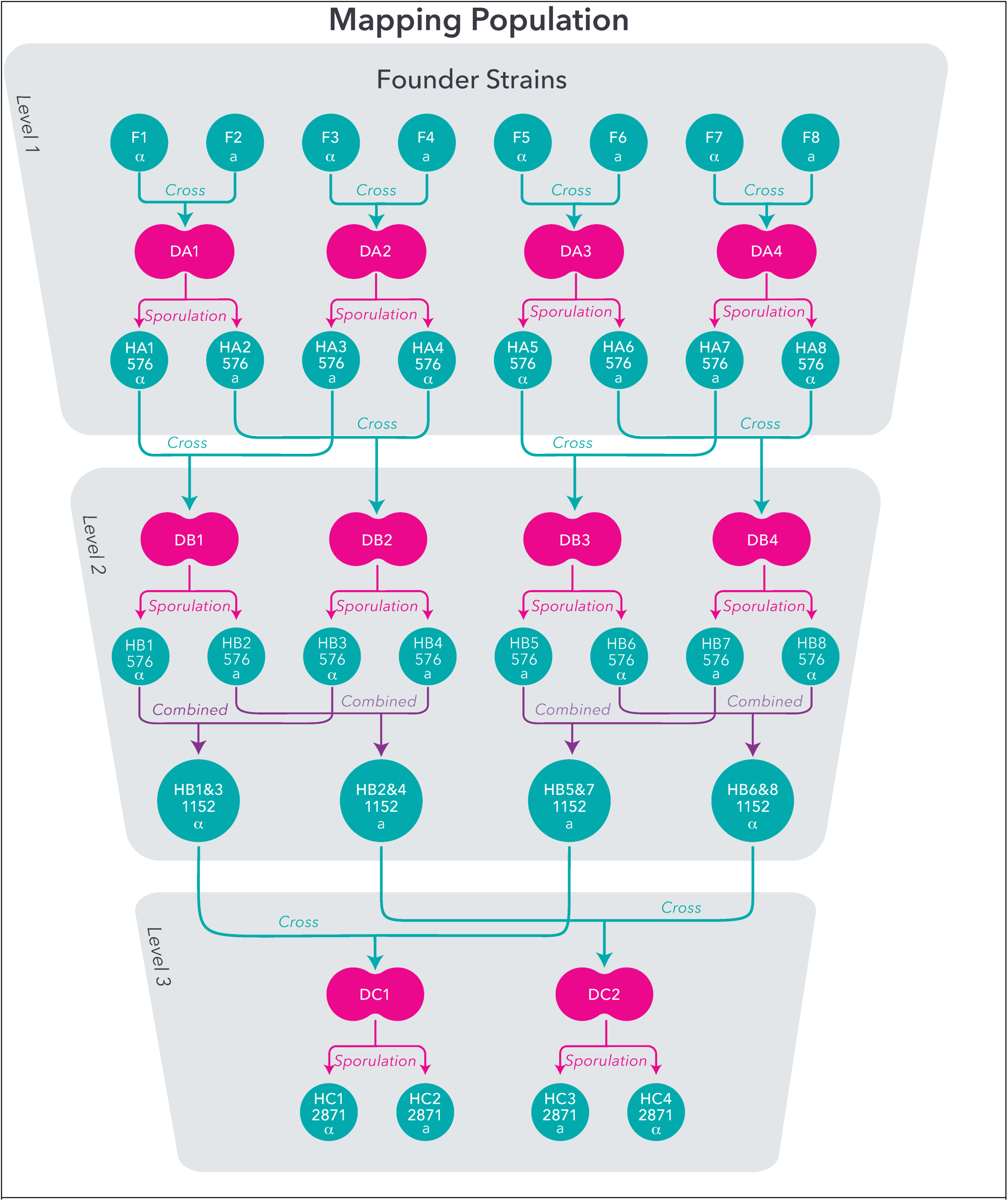
Detailed outline of funnel cross design. Closed shapes refer to populations or founder strains, with red indicating diploids and blue haploids. Blue lines indicate strain or population crossing, red lines indicate sporulation and purple lines indicate populations being combined. The final mapping population consists of the combined populations HC1-4.

**Fig S2.**
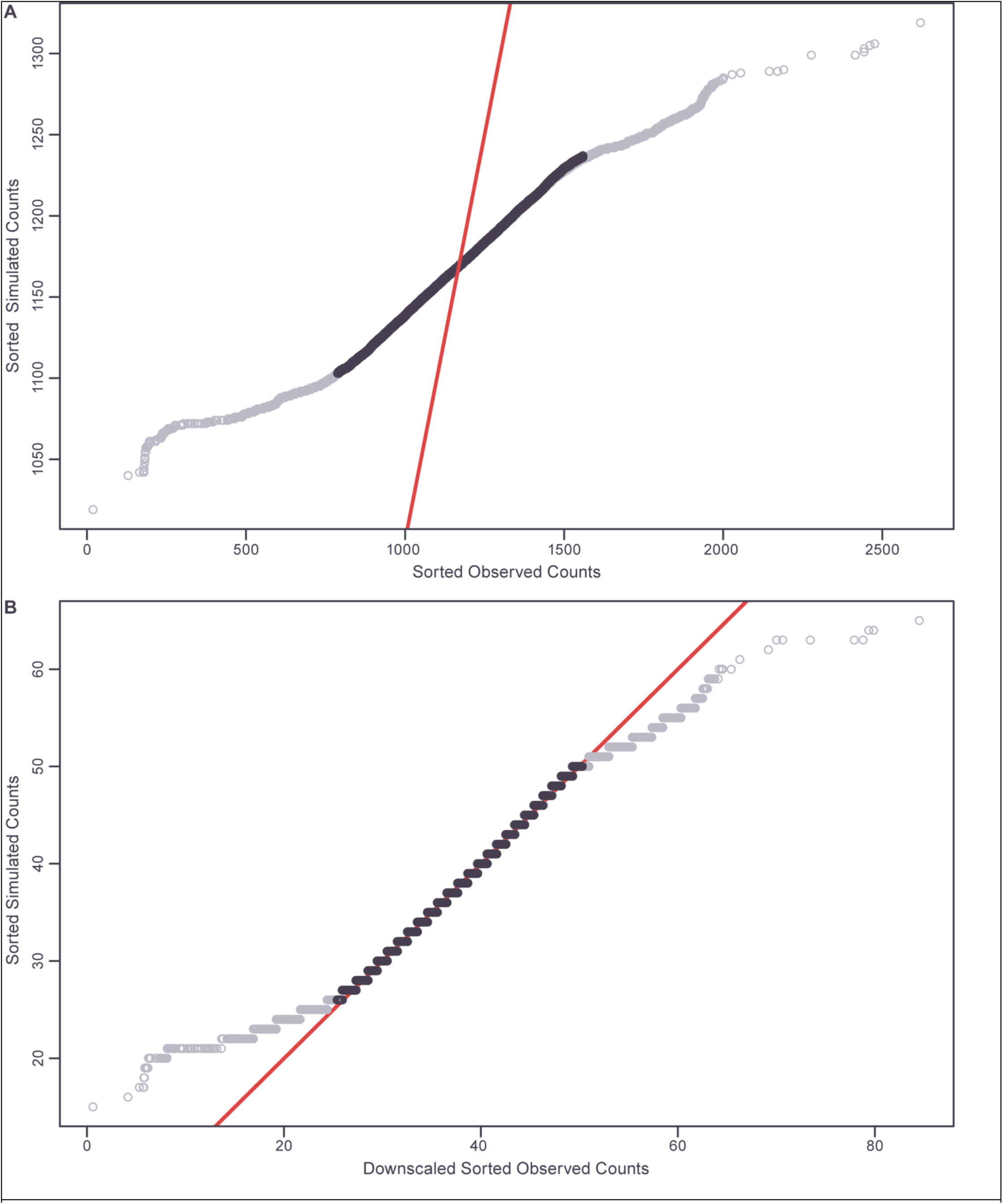
Q-Q plots of observed marker haplotype counts vs simulated Poisson distributions. Observed counts exclude markers on Chromosome IX. Poisson distributions generated using the same sample size and mean as the observed count distributions. Central 95% of values shown in black and x=y line in red. (**A**) Unadjusted haplotype counts. (**B**) Haplotype counts downscaled 31-fold.

**Fig S3.**
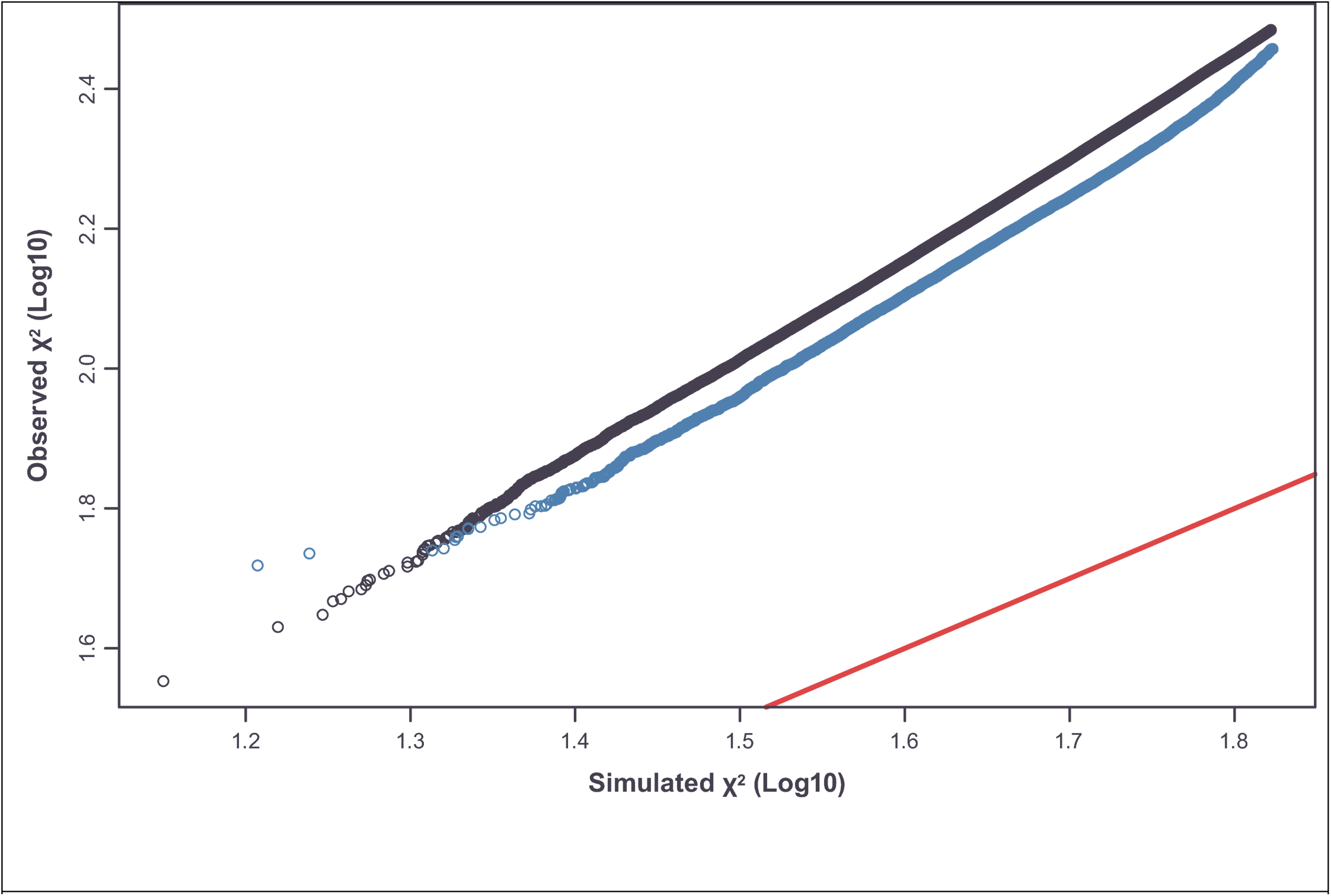
Q-Q plot of interchromosomal pairwise marker scores from the Chi-square test vs simulated Chi-square distribution. 20k values sampled randomly from marker pairs that include Chromosome IX (blue) and 20k values sampled randomly from marker pairs that do not include Chromosome IX (black). Simulated distribution is the same size and has the same degrees of freedom as the observed score distributions. x=y line in red. Upper 5% of scores in each distribution are excluded.

**Fig S4.**
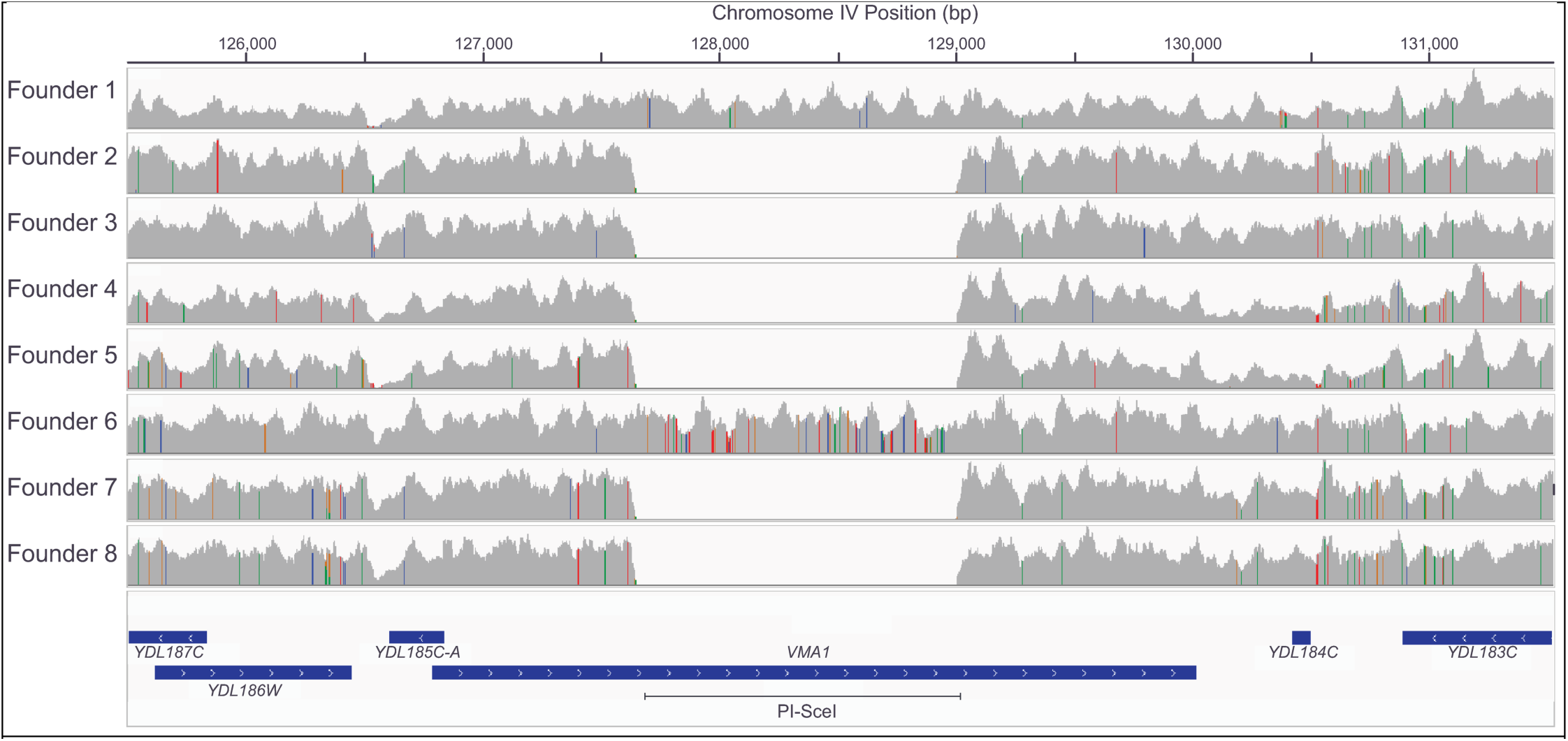
Presence of PI-SceI in founders 1 and 6, but not other founder strains. Visualization of read coverage (relative to the S288c reference genome) across the *VMA1* gene from Illumina sequencing of the founder strains. *VMA1* in the S288c reference genome encodes PI-SceI, with this region within *VMA1* highlighted.

**Fig S5.**
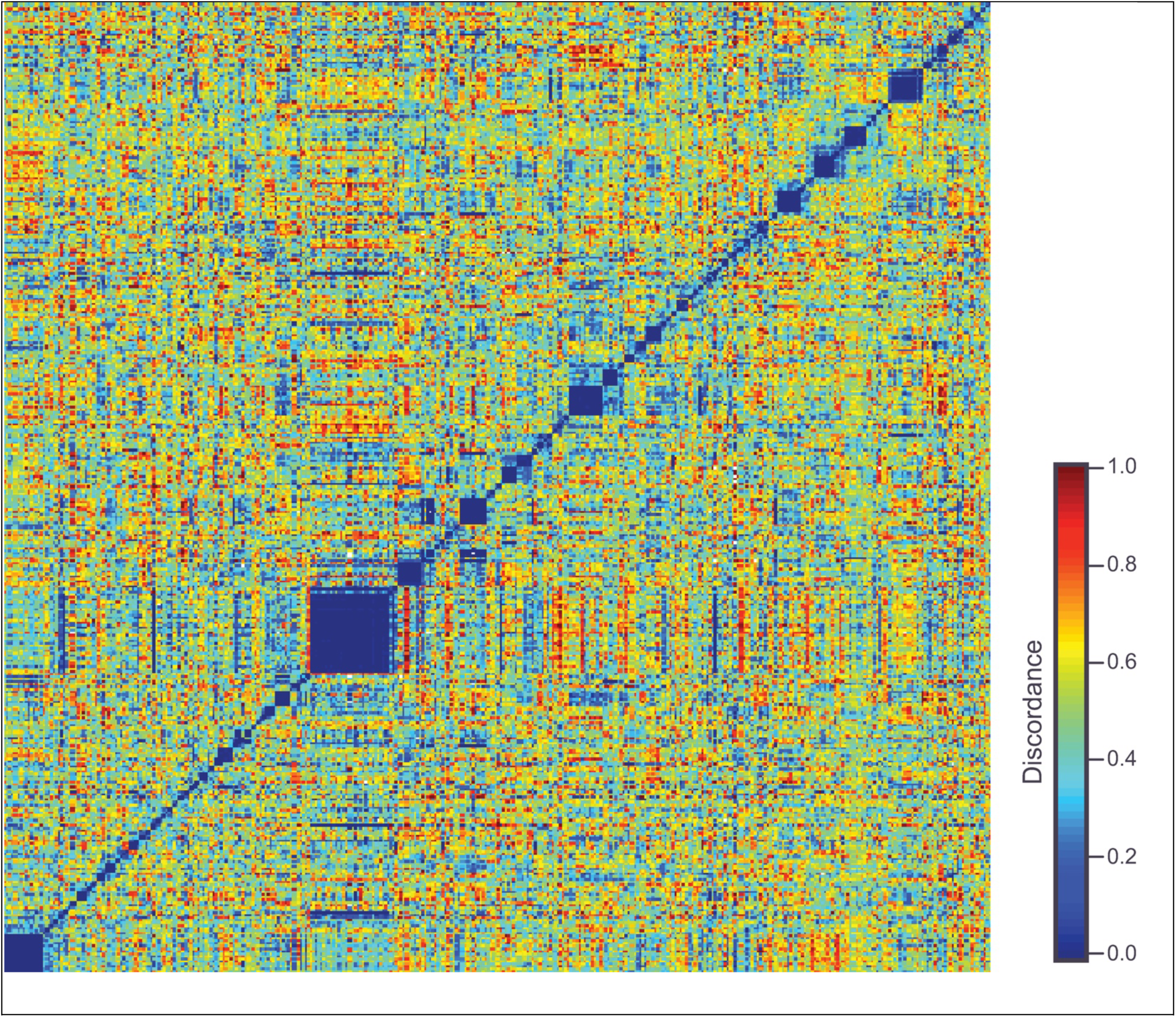
Quantifying shared ancestry among strains disomic for Chromosome. **X**. Strains clustered by pairwise haplotype discordance relative to population G_1_5-6. Distances calculated excluding markers on Chromosomes IX and X. Shared ancestry is expected to produce a discordance value of 0.

## List of Supplementary Files and Tables

**File S1|** Parental alleles at all high-confidence biallelic SNVs used as markers.

**File S2|** Major allele read counts for all euploid strains at all marker positions.

**File S3|** Minor allele read counts for all euploid strains at all marker positions.

**File S4|** Inferred haplotype for each euploid strain at all marker positions.

**File S5|** Subset of markers used in ∼2kb-spaced scaffold.

**File S6|** R data processing script.

**Table S1|** Estimated chromosome copy numbers in each strain.

**Table S2|** Transition probabilities (monosomes) in the final set of mapping strains for any pair of linked makers, as a function of recombinant fraction (R).

**Table S3|** Transition probabilities (disomes) in the final set of mapping strains for any pair of linked makers, as a function of recombinant fraction (R).

